# A human-specific enhancer fine-tunes radial glia potency and corticogenesis

**DOI:** 10.1101/2024.04.10.588953

**Authors:** Jing Liu, Federica Mosti, Hanzhi T. Zhao, Jesus E. Sotelo-Fonseca, Carla F. Escobar-Tomlienovich, Davoneshia Lollis, Camila M. Musso, Yiwei Mao, Abdull J. Massri, Hannah M. Doll, Andre M. Sousa, Gregory A. Wray, Ewoud Schmidt, Debra L. Silver

## Abstract

Humans evolved an extraordinarily expanded and complex cerebral cortex, associated with developmental and gene regulatory modifications^1–3^. Human accelerated regions (HARs) are highly conserved genomic sequences with human-specific nucleotide substitutions. Although there are thousands of annotated HARs, their functional contribution to human-specific cortical development is largely unknown^4,5^. *HARE5* is a HAR transcriptional enhancer of the WNT signaling receptor *Frizzled8 (FZD8)* active during brain development^6^. Here, using genome-edited mouse and primate models, we demonstrate that human *(Hs) HARE5* fine-tunes cortical development and connectivity by controlling the proliferative and neurogenic capacity of neural progenitor cells (NPCs). *Hs-HARE5* knock-in mice have significantly enlarged neocortices containing more neurons. By measuring neural dynamics *in vivo* we show these anatomical features correlate with increased functional independence between cortical regions. To understand the underlying developmental mechanisms, we assess progenitor fate using live imaging, lineage analysis, and single-cell RNA sequencing. This reveals *Hs-HARE5* modifies radial glial progenitor behavior, with increased self-renewal at early developmental stages followed by expanded neurogenic potential. We use genome-edited human and chimpanzee (Pt) NPCs and cortical organoids to assess the relative enhancer activity and function of *Hs-HARE5* and *Pt-HARE5.* Using these orthogonal strategies we show four human-specific variants in *HARE5* drive increased enhancer activity which promotes progenitor proliferation. These findings illustrate how small changes in regulatory DNA can directly impact critical signaling pathways and brain development. Our study uncovers new functions for HARs as key regulatory elements crucial for the expansion and complexity of the human cerebral cortex.

---

Humans are distinguishable from other species with our enlarged cerebral cortex and extraordinary cognitive capacities. Underlying these features is the expansion of neural cell number, diversity and circuitry^2,7^. Relative to other species, humans have a protracted neurogenesis allowing for increased neuron production and maturation^3^. During cortical neurogenesis, the primary neural progenitor cells (NPCs) in the ventricular zone (VZ) are radial glial cells (RGCs). These progenitors undergo self-renewing divisions to expand the precursor pool or differentiating divisions to produce neurons and neurogenic basal progenitors (BPs). In mice, the predominant BPs are intermediate progenitors (IPs). In humans and non-human primates, BPs increase in abundance, composition, and proliferative capacity which amplifies neuronal production^8–10^. Thus, neuron number is collectively defined by the balanced proliferative and neurogenic divisions of neural precursors. Yet, we have a limited understanding of how progenitors are controlled to generate unique features of related species.

Comparative genomics of primates reveal extensive human-specific loci, including chromosomal duplications, deletions, and point mutations^11^. While protein-coding regions are highly conserved between humans and our closest extant relative, the chimpanzee, differences in non-coding DNA are especially prominent and posited to contribute to human-specific features^12^. Human Accelerated Region (HARs) are enriched in non-coding DNA and these highly conserved loci contain single nucleotide changes fixed specifically in humans ^4,5,13–16^. Nearly 3,000 HARs have been identified, many of which are located nearby neurodevelopmental genes, and act as gene regulatory enhancers in neural cells^5,12,14,17–20^. Notably, mutations in HARs are associated with neurological diseases^21,22^. Use of transgenic mice and massively parallel reporter assays *in vitro* have shown HARs have tissue and species-specific enhancer activity regardless of the trans environment^5,13,14^. With two exceptions in skin and the developing limb^23,24^, the direct function of HARs *in vivo* is unknown. Indeed, a key knowledge gap is how the thousands of HARs functionally modify human brain features (Fig.1a).

**Fig. 1.**
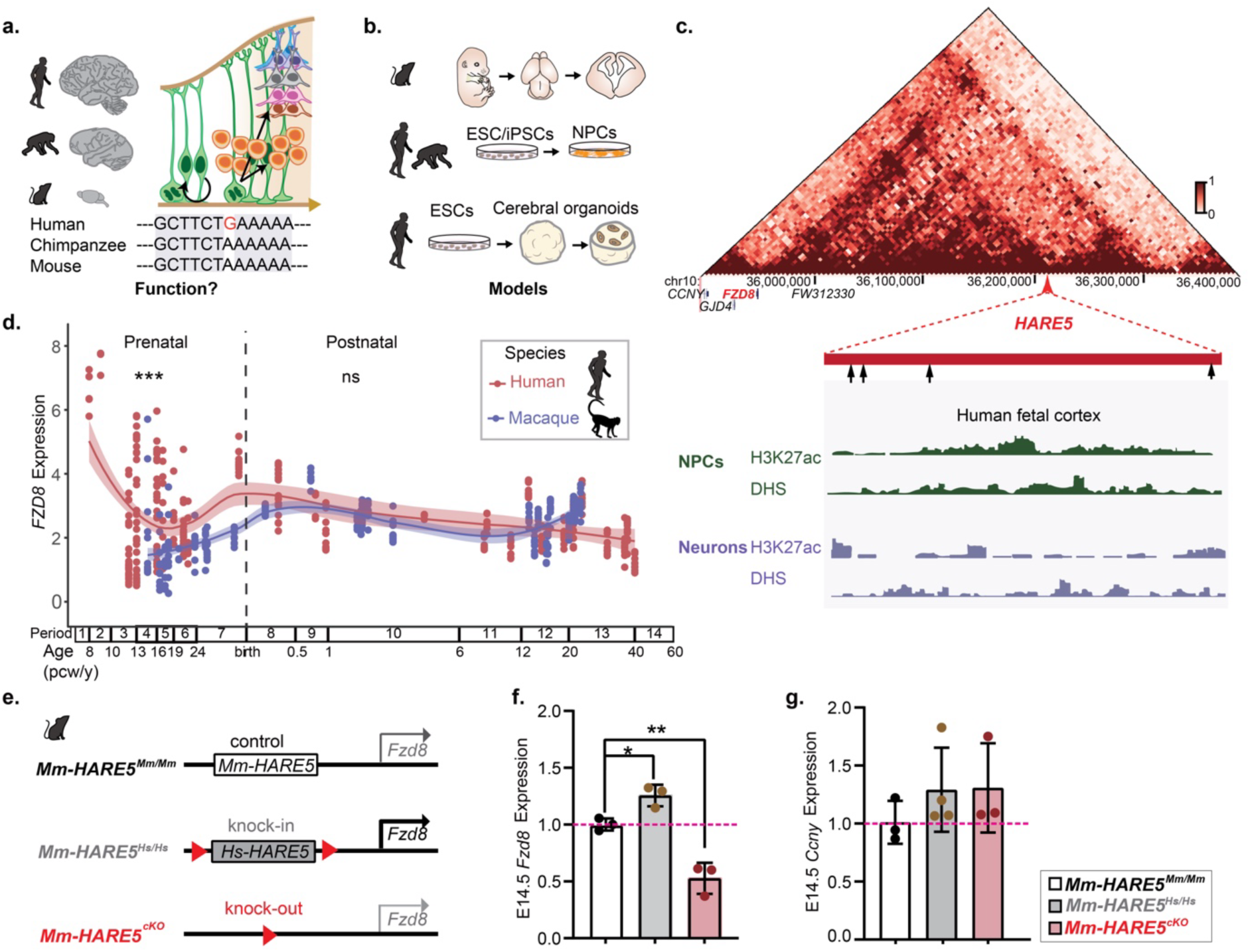
*HARE5* is an active enhancer in developing human neural progenitors. **a**, This study aims to understand the cellular and developmental mechanisms by which human accelerated regions (HARs) regulate brain development? Depicted is an example of a HAR, brains of different species and simplified cartoon of neurogenesis from mice in which radial glial progenitor cells (RGCs, green) can either self-renew, generate neuron-producing intermediate progenitors (IPs, orange), or neurons (blue). **b**, Cartoons of experimental models samples used in this study: mouse embryo and brain (top); 2D neural progenitor cell differentiation from human and chimpanzee ESCs and iPSCs (middle); 3D cortical organoids derived from human ESCs (bottom). **c**, Hi-C map of chromosome 10 region of human fetal brain tissue (top). ChIP-seq data for H3K27ac and DNase hyper-sensitive (DHS) sites for *HARE5* in NPCs and neurons, respectively (bottom). **d**, *FZD8* expression is higher in the human neocortex compared to macaque at prenatal (p<0.001), but not postnatal stages. **e**, *HARE5* enhancer manipulation in mouse models used in this study. **f**, qPCR of *Fzd8* mRNA in E14.5 cortices from control, *HARE5^Hs/Hs^* and *Emx1-Cre*;*HARE5^cKO^*(cKO) mice. Each litter was normalized to wild type littermates. n=3 embryos; 2 litters each genotype. **g**, qPCR of *Ccny* mRNA in E14.5 cortices from control, *HARE5^Hs/Hs^* and *HARE5^cKO^* mice. n=3-4 embryos; 3 litters per genotype. Graphs and bar plot, means ± S.D. *p<0.05, **p<0.01, ***p<0.001. ns, not significant One-way ANOVA with Dunnett’s multiple comparisons test (f, g), Welch’s t-test with Bonferroni correction (d).

We previously discovered *HARE5* (*ANC516*), which diverges between humans and chimpanzees, and shows species-specific enhancer activity in the developing mouse forebrain^6^. *HARE5* is on chromosome 10, with only 4 human-specific nucleotides across 619 bp^5,6^. Chromosome Conformation Capture in the developing mouse cortex showed the endogenous mouse *(Mm) HARE5* is an enhancer of *Fzd8*, a receptor in the WNT signaling pathway implicated in cortical development and disease^25–28^. Transgenic mice (Hs-*HARE5::Fzd8*) in which human *HARE5* (*Hs-HARE5)* drove *Mm-Fzd8* expression, have enlarged cortices^6^. These findings suggest a key role of *Hs-HARE5* in brain development. Yet how this HAR mechanistically controls human-specific features remains unknown. Thus, *HARE5* provides an unprecedented opportunity to explore how human-specific DNA changes in HARs drive brain features (Fig.1a).

Here, we use orthogonal models to discover new mechanisms underlying human-specific cortical development, reliant on modifications of *HARE5* (Fig.1b). Using a knock-in *Hs-HARE5* mouse model we perform in-depth developmental analysis and functional mapping of brain activity. We show that *Hs-HARE5* impacts neuronal number and brain size by modulating proliferative and neurogenic capacity of progenitors and this impacts functional connectivity between cortical regions. Further, using genome-edited human and chimpanzee cells and organoids we discover that human-specific mutations directly promote neurogenesis. These data demonstrate how gene regulatory modifications of a developmental signaling pathway contribute to human-specific brain features.

## *HARE5* activates *FZD8* expression with species divergency

We first characterized *HARE5* activity and target gene expression in the developing human brain. Our previous investigation showed the *HARE5* enhancer is active in the developing mouse and that *Mm-HARE5* is an enhancer of *Fzd8*^6^. To assess *HARE5* enhancer activity in the developing human cerebral cortex, we interrogated published Hi-C, DNaseI-seq and H3K27ac ChIP-seq data^5,29^. This revealed that *HARE5* is an active enhancer in the developing brain and its highest frequency target is *FZD8* (Fig.1c, Extended Data Fig.1a). To validate *HARE5* is a *FZD8* enhancer in human neural cells, we used CRISPR interference (CRISPRi) to target *HARE5* (Extended Data Fig.1b). Analysis of *FZD8* mRNA levels by RT-qPCR showed robust repression after 3 days (Extended Data Fig.1c). This indicates that *HARE5* functions as an active enhancer of *FZD8* in human neural cells.

Given that human and chimpanzee *HARE5* orthologs exhibit differential enhancer activity in mouse embryos^6^, we hypothesized that *FZD8* may likewise show expression divergence between related species. Rhesus macaque is a closely related species to humans and a commonly studied nonhuman primate. We thus probed RNA sequencing datasets of developing human and macaque neocortex from BrainSpan and PsychENCODE^30^. Compared to developing macaque, prenatal human neocortex showed consistently higher *FZD8* expression (Fig.1d). In contrast, in the cerebellum and postnatal cortex, *FZD8* expression levels were comparable between species (Fig.1d, Extended Data Fig.1d). These data indicate human-specific expression patterns of *FZD8* in the developing neocortex.

We next assessed the extent to which *HARE5* activity and *FZD8* expression are cell-type specific in the developing cortex. Notably, *HARE5* showed DNaseI-hypersensitivity (DHS) and H3K27ac epigenetic marks in NPCs but not neurons, reflecting enhancer activity (Fig.1c)^5,29^ Inspection of single cell (sc)RNA-sequencing datasets^31–33^ showed *FZD8* is mainly expressed in cortical RGCs but not newborn excitatory neurons of the developing mouse, macaque and human cortex (Extended Data Fig.1e-g). Thus, while *FZD8* levels are divergent across species, the overarching pattern of expression is conserved. This is consistent with *Hs-HARE5* and *Pt-HARE5* activity patterns previously reported in the mouse^6^. In sum, these data demonstrate that *HARE5* enhancer activity and *FZD8* expression is enriched in NPCs, particularly RGCs, but less detectable in neurons of the developing cerebral cortex.

## *Hs-HARE5* increases cortical size and neuron number in a knock-in mouse

We next tested the hypothesis that *Hs-HARE5* promotes human features in the mouse cortex. Towards this, we generated a knock-in mouse in which both *Mm-HARE5* alleles were replaced with the corresponding human *HARE5 (Mm-HARE5^Hs/Hs^*) (Fig.1e). To assess the ancestral function of *Mm-HARE5* during mouse cortical development, we surrounded this locus with two loxP sites to generate a cKO (*Mm-HARE5^cKO^*). To generate *Mm-HARE5^cKO^* mice, we used *Emx1*-Cre, which is active in cortical progenitors and their progeny beginning at E9.5^34^. For all analyses, we compared these mice with wild-type controls (*Mm-HARE5^Mm/Mm^*) (Fig.1e).

We first assessed *Fzd8* expression in these models by RT-qPCR of the embryonic (E) 14.5 dorsal telencephalon. Compared to *Mm-HARE5^Mm/Mm^* (control) littermates, *Fzd8* mRNA level exhibited a significant, 1.25-fold increase in *Mm-HARE5^Hs/Hs^*. In contrast, *Fzd8* showed a 50% decrease in the *Mm-HARE5^cKO^* cortex (Fig.1f). As a control, we performed RT-qPCR for *Ccny*, which is located in the same chromosomal region, but is a lower frequency *HARE5* target gene from human NPC HiC data (Extended Data Fig.1a)^29^. Neither *Mm-HARE5^Hs/Hs^* nor *Mm-HARE5^cKO^* significantly impacted *Ccny* levels in the developing mouse brain (Fig.1g). Collectively, these data demonstrate *Hs*-*HARE5* is an enhancer of *Fzd8* with stronger activity than *Mm-HARE5* in the developing mouse cortex.

We next evaluated the impact of *Hs-HARE5* upon brain size. Cortical area quantification of 3 month postnatal brains showed a significant increase of *Mm-HARE5^Hs/Hs^* compared to control littermates (Fig.2a,b). Notably, both heterozygote (*Mm-HARE5^Hs/Mm^*) and homozygote (*Mm-HARE5^Hs/Hs^*) showed the same trend upon cortical size at P0 (Extended Data Fig.2a,b). In contrast to this phenotype, loss of either one or both copies of *Mm-HARE5* from neural progenitors and their progeny led to profound microcephaly, with a 5-10% cortical area decrease compared with control littermates (Extended Data Fig.2a,b). These data demonstrate that the ancestral mouse *HARE5* enhancer is essential for cortical size, whereas human *HARE5*, due to stronger activity, has the opposite impact to increase cortical size.

**Fig. 2.**
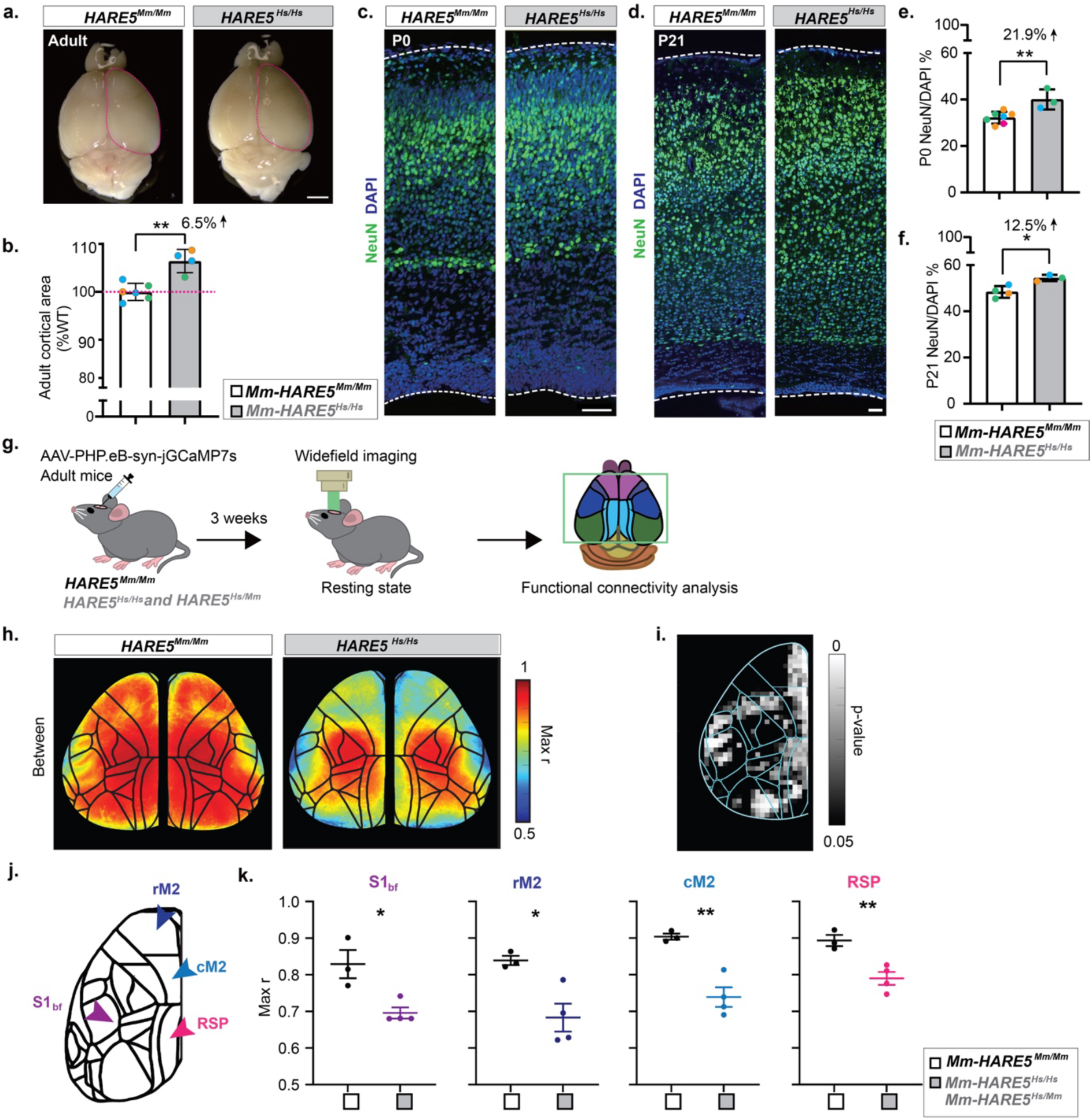
*Hs-HARE5* promotes cortical expansion and neocortical connectivity changes in mice. **a**, Representative whole mount images of control and *HARE5^Hs/Hs^* brains at 3 months. Dotted line is control superimposed on both brains. **b**, Quantification of cortical area in 3 month brains. n=4-6 animals; 3 litters per genotype. Each litter normalized with wildtype littermates. **c**, Representative sections of P0 control and *HARE5^Hs/Hs^*brains stained with NeuN (green) and DAPI (blue). **d**, Representative sections of P21 control and *HARE5^Hs/Hs^*brains stained with NeuN (green) and DAPI (blue). **e**, Quantification of NeuN+ cells in P0 control and *HARE5^Hs/Hs^*cortices. n=3-7 animals; 4 litters per genotype. **f**, Quantification of NeuN+ cells in P21 control and *HARE5 ^Hs/Hs^* cortices. n=3-4 animals; 3 litters per genotype. **g**, Experimental paradigm for functional mapping of adult mice. **h**, Widefield calcium imaging of resting state activity across the dorsal cortex shows lower inter-region correlation in *HARE5* knock-in mice. Average maps show the pixelwise correlation value threshold of independence. Each pixel’s activity only correlates with one functional cluster above this threshold. *n*=3 WT and *n*=4 *HARE5* knock-ins (n=3 *Mm-HARE5^Hs/Hs^*, n=1 *Mm-HARE5^Hs/Mm^*). **i**, Correlation maps were spatially binned and compared between genotypes using two-tailed t-tests (All non-black pixels are statistically significant, p<0.05). **j**, The Allen Atlas Common Coordinate Framework was used to group functional clusters belonging to the same known anatomical regions (*left*). Motor regions (M1 and M2) were split rostral-caudally and evaluated separately. **k**, Graphs show average maximum Pearson’s correlation (r) values of each region per genotype, where each data point represents an animal (*right*). Scale bars: 2mm (g), 50µm (i, j). Graphs and bar plot, means ± S.D. *p<0.05, **p<0.01, ***p<0.001. ns, not significant. Student’s unpaired, two-tailed t-test (b, e, f, k).

We next tested if larger brain size was associated with more neurons. We performed NeuN staining on P0 and P21 mouse cortices (Fig. 2c,d). Quantification of NeuN positive cells revealed significantly more mature neurons in the *Mm-HARE5^Hs/Hs^* mouse cortex compared to controls at both stages (Fig. 2e,f). We assessed if these neurons showed altered organization by quantifying the density of NeuN positive cells within cortical columns. In *Mm-HARE5^Hs/Hs^* there were significantly more NeuN cells in upper layer bins at P21 (bins 6, 8-10) while P0 showed no significant change (Extended Data Fig.2c,d). Together, these data demonstrate that *Hs-HARE5* promotes cortical expansion and neuron number in the mouse neocortex.

## *Hs-HARE5* impacts functional connectivity in the cortex

We next aimed to understand if this increased enhancer activity, cortical size and neuron number impacted functional activity across the cortex. A previous study in which more cortical neurons were chemically induced showed this increased neuron ensemble number and functional segregation of local neural networks within the visual cortex^35^. We predicted that cortical expansion and increased neuron number in *Hs-HARE5* mice may similarly alter baseline functional network dynamics. To test this, we mapped resting state dynamics across the dorsal cortex using widefield calcium imaging in awake mice. Neural dynamics during resting state provide insight into the functional connectivity between cortical regions (Fig.2g) ^36,37^. We performed spatiotemporal clustering to parcellate the cortex into functional clusters. On a pixelwise basis, we calculated the threshold correlation value at which each pixel is no longer correlated with any cluster other than its own, as a measure of inter-region independence. Overall, both homozygous (*Mm-HARE5^Hs/Hs^*) and heterozygous (*Mm-HARE5^Hs/Mm^*) mice exhibited more functionally independent activity compared to control (Fig.2h and Extended Data Fig.3a), and significantly so in both primary sensory regions (whisker barrel field, S1_bf_), as well as premotor (rostral and caudal M2) and higher order associative regions (retrosplenial, RSP) (Fig.2i-k). We also examined intra-region correlation as a measure for how well each pixel correlates with its own functional cluster. As with our analysis of correlation between cortical regions, *Hs-HARE5* mice exhibited lower correlation within clusters (Extended Data Fig.3b-e). Taken together, these data show that *Hs-HARE5* alters the functional properties of cortical networks by increasing functional independence between cortical regions.

## *Hs-HARE5* amplifies neural progenitors at early stages of cortical development

We next investigated the developmental basis for *Hs-HARE5*-mediated postnatal brain features. To better understand which cells display *Hs-HARE5* activity across different stages of the developing mouse cortex, we took advantage of a previously generated mouse, *Hs-HARE5*::EGFP-PEST^6^, in which EGFP is a reporter of enhancer activity (Fig.3a). *HARE5* enhancer activity was remarkably high in the developing neocortex at E11.5 and E14.5, mainly within the germinal regions (Fig.3a). To assess cell-specific activity in E14.5 RGCs and IPs, we co-stained sections against PAX6 and TBR2, respectively (Fig. 3b). While active in a small fraction of IPs, *Hs-HARE5* was especially active in about 50% of RGCs (Fig.3c; Extended Data Fig.4a). These results indicate that *Hs-HARE5* drives robust expression in radial glia of the developing telencephalon. This observation is consistent with the active epigenetic marks of *HARE5* specifically in human NPCs (Fig.1c), and the progenitor-specific expression of *Fzd8* in mice and primate brains (Extended Data Fig.1e-g).

**Fig. 3.**
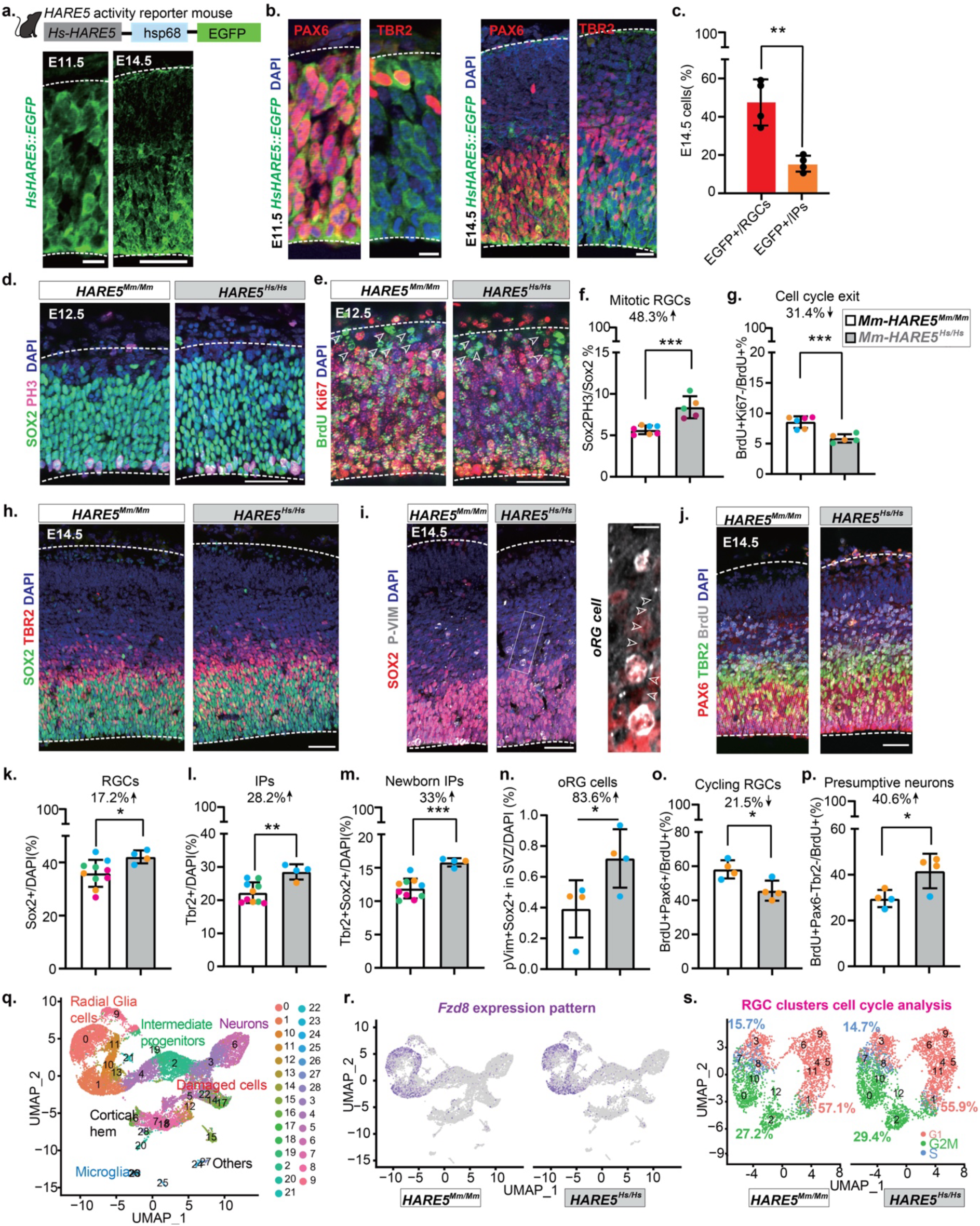
*Hs-HARE5* promotes neural progenitor expansion during cortical development. **a**, Schematic of *Hs-HARE5*::EGFP reporter mouse line. Representative sections of E11.5 and E14.5 *Hs-HARE5*::EGFP brains indicating *HARE5* enhancer activity by EGFP. **b**, Representative sections of E11.5 and E14.5 *Hs-HARE5*::EGFP brains stained with PAX6 (red-left), TBR2 (red-right) and DAPI (blue). **c**, Quantification of proportion of RGCs (PAX6+ or SOX2+) and IPs (TBR2+) that are EGFP+ in E14.5 *Hs-HARE5*::EGFP cortices. n=4 embryos. 3 litters. **d**, Representative sections of E12.5 control and *HARE5^Hs/Hs^* brains stained with SOX2 (green), PH3 (magenta) and DAPI (blue). **e**, Representative sections of E12.5 control, *HARE5^Hs/Hs^* brains stained with BrdU (green), Ki67 (red) and DAPI (blue). **f**, Quantification of mitotic RGCs in E12.5 control and *HARE5^Hs/Hs^* cortices. n=3-7 embryos; 4 litters each genotype. **g**, Quantification of cell cycle exit in E12.5 control, *HARE5^Hs/Hs^* cortices. n=3-6 embryos from 4 different litters per condition. **h**, Representative sections of E14.5 control and *HARE5^Hs/Hs^* brains stained with SOX2 (green), TBR2 (red) and DAPI (blue). **i**, Representative sections of E14.5 control and *HARE5^Hs/Hs^*brains stained with SOX2 (red), P-VIM (grays) and DAPI (blue). High magnification view of oRG cells (arrowheads depict the basal process). **j**, Representative sections of E14.5 control and *HARE5^Hs/Hs^* brains stained with PAX6 (red), TBR2 (Green), BrdU (grays) and DAPI (blue). **k-n,** Quantification of RGCs (**k**), IPs (**l**), newborn IPs (**m**), oRGs (**n**), cell cycle exit (**o**), and presumptive newborn neurons (**p)** in E14.5 control and *HARE5 ^Hs/Hs^* cortices. n=4-10 embryos; 2-4 litters per genotype. **q**, UMAP of sc-RNAseq from E14.5 control and *HARE5 ^Hs/Hs^* cortices. n=2 embryos; 2 litters each genotype. **r**, UMAP showing *Fzd8* expression pattern. **s**, UMAP of cell cycle phase of RGCs. Scale bars: 10µm (a left, i right), 50µm (a right), 50µm (b, d, f, h, i left, j). Graphs and bar plots, means ± S.D. *p<0.05, **p<0.01, ***p<0.001. Student’s unpaired, two-tailed t-test (c, f, g, k-p).

As *Mm-HARE5^Hs/Hs^* mouse brains are enlarged with more cortical neurons (Fig.2c-f), we predicted that *Hs-HARE5* impacts neurons by acting in neural progenitors. To investigate this possibility, we first assessed progenitors at the onset of neurogenesis when neuroepithelial cells (NEs) have transitioned into RGCs. At E12.5, the majority of RGCs undergo proliferative divisions to expand the progenitor pool while some begin to produce neurons. The overall number of SOX2+ RGCs was unaffected in the *Mm-HARE5^Hs/Hs^* cortex (Extended Data Fig.4f). However, we observed a striking increase in mitotic RGCs in *Mm-HARE5^Hs/Hs^* cortices (Fig.3d,f). The *Mm-HARE5^Hs/Hs^* progenitors exhibited reduced cell cycle exit, indicating that at E12.5 they undergo increased self-renewal (Fig.3e,g). These findings corroborate previous observations of the *Hs-HARE::Fzd8* transgenic mouse^6^. Altogether these data demonstrate that *Hs-HARE5* promotes proliferative capacity and self-renewal of early stage RGCs.

Given that *Hs-HARE5* promotes RGC mitosis and self-renewal at E12.5, we asked whether ancestral *Mm-HARE5* impacts neural progenitors. In contrast to the *Hs-HARE5* knock-in cortices, *Mm-HARE5 ^cKO^* cortices had significantly fewer RGCs compared to control, with no impact on mitotic index (Extended Data Fig.4b,c,f). Also in contrast to the knock-in, *Mm-HARE5 ^cKO^* RGCs showed increased cell cycle exit at this stage (Extended Data Fig.4d,e). Taken together, these findings indicate that *HARE5* is necessary and sufficient for RGC expansion at the onset of neurogenesis.

## *Hs-HARE5* promotes neural differentiation at mid-cortical development

Over the course of development, the relative proportion of RGCs decreases as they produce more neurons and IPs^38^. This increase in abundance and complexity of IPs is thought to contribute to neocortical enlargement in primates^2,7^. Therefore, we assessed whether *Hs-HARE5* impacts progenitor composition at peak stages of neurogenesis. We quantified RGCs and IPs by staining for SOX2 and TBR2, respectively, in E14.5 cortices (Fig.3h,i). Notably, RGCs were significantly increased by 17.2% in the *Mm-HARE5^Hs/Hs^* cortex compared to control (Fig.3k). The IP population also showed a striking 28.2% increase in *Mm-HARE5^Hs/Hs^* cortices (Fig.3l). We also observed significantly more newborn IPs (TBR2+SOX2+) in *Mm-HARE5^Hs/Hs^*brains (Fig.3m). Notably, both heterozygote (*Mm-HARE5^Hs/Mm^*) and homozygote (*Mm-HARE5^Hs/Hs^)* mice showed the same trends for both progenitors (Extended Data Fig.4j,k). We also assessed oRGs, a major BP of primates and gyrencephalic species. For this we quantified cells expressing SOX2 and Phospho-vimentin and exhibiting a unipolar morphology and basal location in the SVZ (Fig.3i). This showed a slight but significant increase in oRGs in the E14.5 *Mm-HARE5^Hs/Hs^* cortex (Fig.3n). Consistent with these data, mitotic cells in *Mm-HARE^Hs/Mm^* cortices were enriched basally, suggesting the possibility that basal progenitors may undergo increased proliferation (Extended Data Fig.4g,h). Altogether, this demonstrates that at mid-stages of neurogenesis, *Hs-HARE5* promotes more RGCs and basal progenitors, including both IPs and oRGs.

Given the increase in both RGCs and newborn IPs, we hypothesized that at E14.5 *Mm-HARE5^Hs/Hs^* progenitors may undergo increased production of neurons. To assay this, we measured cell cycle exit of both IPs and RGCs using BrdU labeling at E13.5, with subsequent staining of TBR2, PAX6 and BrdU at E14.5 (Fig.3j). Compared to controls, *Mm-HARE5^Hs/Hs^*brains contained significantly fewer BrdU+PAX6+ cells, indicating a higher proportion of RGCs exit the cell cycle (Fig.3o). In comparison, there was no notable change in cell cycle exit of IPs (Extended Data Fig.4i). Additionally, we quantified significantly more BrdU+Sox2-Tbr2-cells (presumed post-mitotic neurons) in *Mm-HARE5^Hs/Hs^* cortex compared to control (Fig.3p), consistent with increased neural differentiation. These data demonstrate that *Hs-HARE5* promotes RGC cell cycle exit at E14.5, resulting in production of more excitatory neurons.

Collectively, analyses at these two developmental stages reveal that *Hs-HARE5* modulates progenitor behavior over the course of cortical development. *Hs-HARE5* promotes proliferation and self-renewal of RGCs at the onset of neurogenesis and continues to expand the precursor pool while amplifying neuronal and progenitor generation as development proceeds.

## scRNA sequencing reflects neural progenitor cell state and composition changes

To further probe how *Hs-HARE5* influences cellular composition we used single cell sequencing. Towards this, we performed 10x genomics single-cell RNA sequencing (scRNA-Seq) on E14.5 dorsal telencephalons of *Mm-HARE5^Hs/Hs^* and littermate *Mm-HARE5*^Mm/Mm^ (Extended Data Fig.5a). Louvain clustering revealed 28 clusters in both control and *Mm-HARE5^Hs/Hs^* brains. We assigned cell types to distinct clusters based on their marker gene expression and identified RGCs, IPs, neurons, and non-neural cells (Fig.3q). In control brains, *Fzd8* expression was primarily limited to RGCs, consistent with scRNA-Seq datasets of mice and primates^31–33^ (Fig.3r; Extended Data Fig.1e-g). This cell-specific pattern held true in *HARE5^Hs/Hs^*brains, with overall elevated *Fzd8* levels within RGCs (Fig.3r; Extended Data Fig.5b). Given the robust expression of *Fzd8* in RGCs, we focused on these progenitors. Amongst 12 RGC subclusters, the composition differed between control and *Mm-HARE5^Hs/Hs^* suggesting *Hs-HARE5* may impact cell fate or state of RGCs (Extended Data Fig.5c,d). Indeed, the proportion of G2/M RGCs was increased relative to other cell cycle phases in the *Mm-HARE5^Hs/Hs^*cortex (Fig.3s). As *Mm-HARE5^Hs/Hs^* brains also contained more IPs at E14.5 (Fig.3k-m), we assessed the state of IPs. Towards this, we focused only on clusters with high *Tbr2* expression (Extended Data Fig.5e). The subcellular composition among these 14 subclusters changed between control and *Mm-HARE5^Hs/Hs^* (Extended Data Fig.5f). Consistent with observations of RGCs, cell cycle analysis of IPs revealed enriched G2/M populations in the *Mm-HARE5^Hs/Hs^* cortex (Extended Data Fig.5g,h). Taken together, these scRNA-seq data provide orthogonal evidence that *Hs-HARE5* promotes expression of *Fzd8* in RGCs, with changes in cell cycle composition of both RGCs and IPs.

## Human *HARE5* impacts RGC progeny cell fate

From our immunofluorescence analysis and scRNA sequencing, we conclude that *Hs-HARE5* acts in RGCs to initially promote their expansion and subsequently increase genesis of IPs and neurons. This suggests that *Hs-HARE5* may modulate the types of divisions which RGCs undergo. RGCs undergo self-renewing divisions, IP generating divisions, or neurogenic divisions (Fig.4c). We therefore employed live imaging to quantify RGC divisions and progeny generation. We used *Tbr2*-EGFP to monitor production of IPs as described previously (Fig.4a)^39,40^. At E14.5, RGCs were identified as EGFP negative dividing cells. After 20h of imaging primary mouse progenitors, we monitored direct progeny by immunostaining for RGCs (TUJ1-EGFP-), IPs (EGFP+TUJ1-), and neurons (TUJ1+) (Fig.4a-c). Most control RGCs underwent self-renewing divisions, with additional RGCs undergoing differentiating divisions to produce either IPs or Neurons (Fig.4c, d). This is consistent with previous imaging at this stage^39,41^. In contrast, *Mm-HARE5^Hs/Mm^* RGCs underwent significantly more IP producing divisions, but no significant change in self-renewing or neurogenic divisions (Fig.4d). These data demonstrate that *Hs-HARE5* causes RGCs to directly produce more IPs, which is also consistent with more newborn IPs (Fig.3m).

**Fig. 4.**
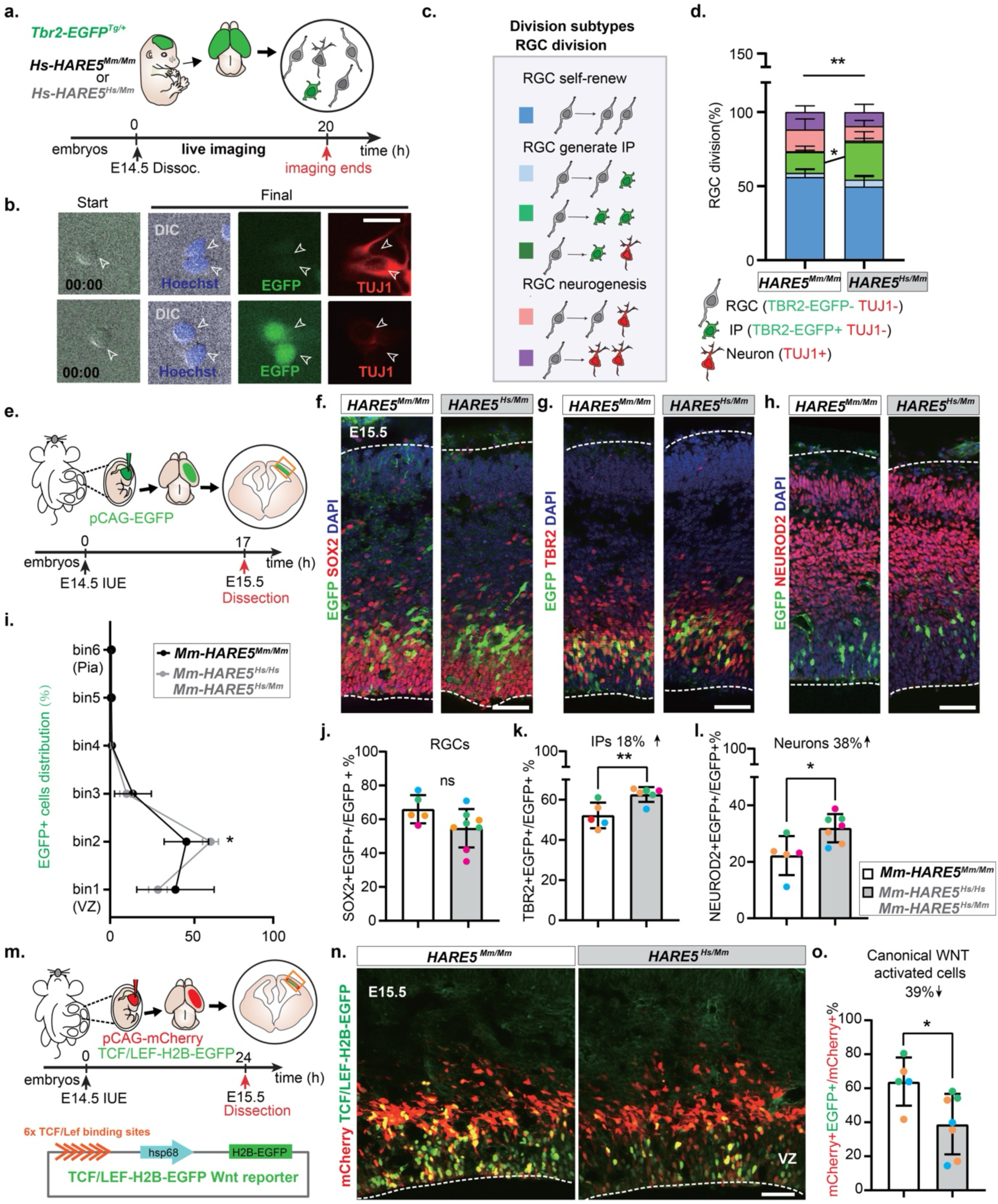
*Hs-HARE5* causes RGC to increase production of neurons and IPs. **a**, Overview of experiment for live imaging primary cortical progenitors and visualizing cell fate using *Tbr2*-EGFP^Tg/+^ (IPs, green). **b**, Representative images to show the different division modes. (Top) RGC symmetric neurogenic division, (bottom) RGC symmetric division making IPs. Tuj1 (red) and Hoechst (blue). **c**, Cartoon shows overview of all division subtypes of RGCs. **d**, Quantification of proportion of control and *HARE5^Hs/Hs^* RGC divisions. n=300 (control) and 420 (*HARE5^Hs/Mm^*) total divisions from 3 independent experiments. **e**, Overview of experiment for lineage tracing by IUE. **f-h,** Representative sections of E15.5 control and *HARE5^Hs/Mm^* brains stained with SOX2 (red) and DAPI (blue) (**f);** TBR2 (red) and DAPI (blue) (**g**); NEUROD2 (red) and DAPI (blue) (**h**). **i**, Binned quantification of EGFP+ distribution in E15.5 control and *HARE5^Hs/Mm^* cortices. Bin 1 is in the ventricular zone (VZ), and Bin 6 is at the pia. X-axis represents the percentage. n=4-5 embryos; 3 litters per genotype. **j-l**, Quantification of generation of RGCs (**j**), IPs (**k**), and neurons (**l**) in E15.5 control and *HARE5* knock in cortices. n=4-10 embryos; 4 litters per genotype. **m**, Overview of experiment for WNT canonical signaling reporter (TCF1/LEF-H2B-EGFP) introduced by IUE. **n**, Representative sections of E15.5 control and *HARE5^Hs/Mm^* brains electroporated with pCAG-mCherry (red) and TCF1/LEF-H2B-EGFP reporter (green). **o**, Quantification of EGFP+ proportion in E15.5 control and *HARE5* knock in cortices. n=5-7 embryos; 3 litters per genotype. Scale bars: 50µm (f, g, h, n), 20µm (b). Graphs and bar plots, means ± S.D. *p<0.05, **p<0.01. Chi-square test with two-way ANOVA Bonferroni’s multiple comparisons test (d), student’s unpaired, two-tailed t-test (j, k, l, o), two-way ANOVA with Bonferroni’s multiple comparisons test (i, d-individual division types).

We next assessed how *Hs-HARE5* impacts RGC progeny *in vivo.* For this we performed *in utero* electroporation (IUE) to sparsely label dividing RGCs for lineage tracing (Fig.4e). In contrast to control brains, *Hs*-*HARE5* led to significantly more EGFP positive cells in the SVZ (Fig.4i). We quantified the identify of these EGFP+ progeny using SOX2, TBR2 and NEUROD2 staining (Fig.4f-h). *Hs-HARE5* RGCs produced significantly more IPs and neurons, but not RGCs, compared to control (Fig.4j-l). This is consistent with the increased cell cycle exit of E14.5 *Hs-HARE5* RGCs (Fig.3o,p). Altogether, these lineage tracing experiments demonstrate that *Hs-HARE5* promotes RGC differentiation into IPs and newborn neurons at E14.5.

To better understand the molecular mechanisms driving the cell fate transition from early to mid-neurogenesis, we performed transcriptome analysis of E12.5 and E14.5 *Mm-HARE5^Mm/Mm^* and *Mm-HARE5^Hs/Hs^*cortices (Extended Data Fig.6a,b). Examination of differentially expressed genes changes across E12.5 to E14.5 revealed 2928 genes in common between control and *HARE5^Hs/Hs^* (Extended Data Fig.6c). Gene Ontology (GO) analysis of these shared genes revealed a significant enrichment in transcripts associated with neuronal differentiation and maturation (Extended Data Fig.6c, d). Expression of 1645 transcripts were specifically changed in the *HARE5^Hs/Hs^*cortex from E12.5 to E14.5, with 870 downregulated and 771 upregulated (Extended Data Fig.6c). The downregulated transcripts were significantly enriched for genes associated with cell cycle and mitosis, whereas the upregulated transcripts were enriched for neural differentiation genes (Extended Data Fig.6e,f). These molecular changes are consistent with observations of the developmental impact of *Hs-HARE5* upon RGCs, with reduced cell cycle exit at E12.5 and increased neural differentiation at E14.5.

We further probed how *HARE5* may impact the WNT pathway, as Frizzled receptors transduce WNT signals^42,43^. As the target gene of *HARE5*, *FZD8* can mediate both canonical and non-canonical signaling. To measure the extent to which *Hs-HARE5* impacts WNT signaling *in vivo*, we performed IUE at E14.5 with a TCF1/Lef-H2B-EGFP canonical WNT reporter^44^, combined with pCAG-mCherry to demarcate electroporated cells (Fig. 4m). The sensitivity of this reporter was validated using a canonical WNT inhibitor IWR1-treated HEK cells (Extended Data Fig.7a, b). Notably, within EGFP+mCherry+ transfected cells canonical WNT signaling was significantly decreased in the E14.5 *Hs-HARE5* cortex (Fig.4n-o). This suggests that increased neural differentiation by *Hs-HARE5* is associated with reduced canonical WNT signaling. These findings are consistent with studies showing that during *in vitro* neural differentiation of human NPCs and *in vivo* mouse cortical neurogenesis, canonical WNT signaling decreases^26,28,45,46^.

## Increased *HARE5* enhancer activity relies on 4 human-specific mutations

Our data using mouse models indicate that *Hs-HARE5* activates RGC expansion and neuron production. A key question is whether this is also true in human NPCs. We first investigated *Hs-HARE5* enhancer activity in human NPCs. *Hs-HARE5*, *Pan troglodytes (Pt)- HARE5* and *Mm-HARE5* nanoluciferase constructs were nucleofected into 2D differentiated forebrain cortical NPCs at day (D)10 along with Fluc (firefly luciferase vector). Luciferase levels were detected 24h later (Fig.5a,b). As expected, *Hs-HARE5* was active in human NPCs with significantly higher activity than both *Pt-HARE5* and *Mm-HARE5* (Fig.5d). This parallels species-specific activity evident in mouse models^6^.

**Fig. 5.**
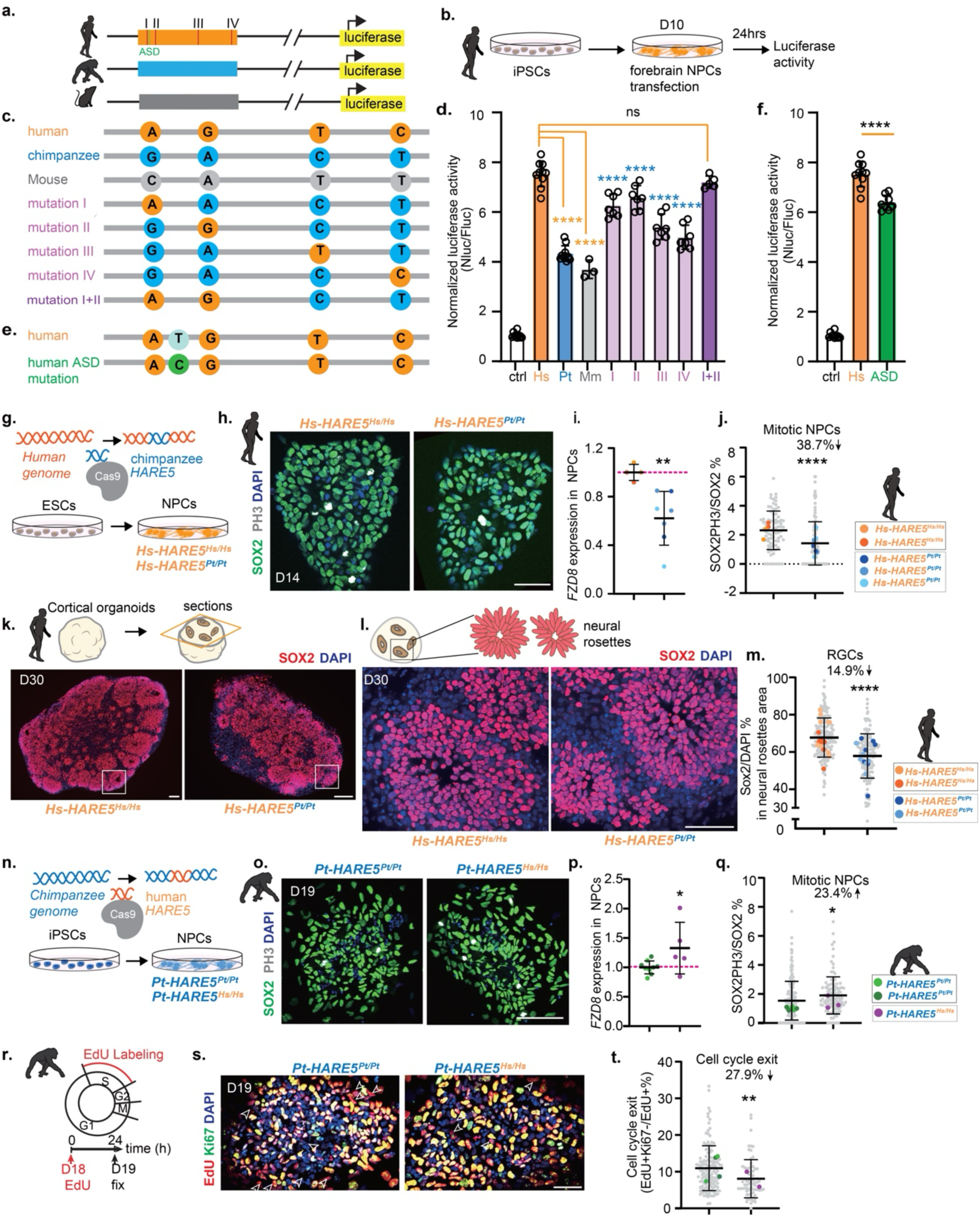
Human-specific variants in *HARE5* directly impact human and chimpanzee NPC proliferation by altering enhancer activity. **a,b**, Experimental paradigm for testing enhancer activity of *Hs-HARE5* mutations by luciferase in human NPCs. **c**, Cartoon shows different mutations generated for luciferase experiments. **d**, Quantification of enhancer activity. n=3 individual differentiations; dots represent wells. **e**, Cartoon shows ASD mutation generated for luciferase experiments. **f**, Quantification of enhancer activity. n=3 individual differentiations; dots represent wells. **g**, Experimental paradigm for generating *HARE5^Pt/Pt^*knock-in cell lines using H9 cells and differentiated into NPCs. **h**, Representative image of D14 control and *HARE5^Pt/Pt^* human NPCs stained with SOX2 (green), PH3 (grays) and DAPI (blue). **i**, qPCR of *FZD8* mRNA in D12-14 NPCs from control, *HARE5^Pt/Pt^* human NPCs. n=2-3 cell lines each condition (represented by color); 2-3 individual differentiations shown as individual dots each. **j**, Quantification of mitotic RGCs of D14-D16 control and *HARE5^Pt/Pt^*human NPCs. n=2-3 cell lines of each condition; 2-3 individual differentiations; grey dots represents an NPC island imaged and quantified; color-coded dots represents the average value of the cell line from each differentiation. **k**, Representative sections of D30 control and *Hs-HARE5^Pt/Pt^* human cortical organoids stained with SOX2 (red) and DAPI (blue). **l**, Representative sections of D30 control and *HARE5 ^Pt/PT^* human cortical organoid depicting neural rosettes regions stained with SOX2 (red) and DAPI (blue). **m**, Quantification of RGCs (SOX2+) of D30 control and *Hs-HARE5^Pt/Pt^* human cortical organoids within neural rosette regions. n=2 cell lines of each condition; 2 individual differentiation; grey dot represents a neural rosettes region imaged and quantified; each color-coded dot represents the averaged value of the single organoid from each differentiation. **n**, Experimental paradigm for generating *HARE5^Hs/Hs^* knock-in cell line using C3649 iPSCs and differentiated into chimpanzee NPCs. **o**, Representative image of D19 control and *HARE5^Hs/Hs^*chimp NPCs stained with SOX2 (green), PH3 (grays) and DAPI (blue). **p**, qPCR of *FZD8* mRNA in D15 control and *Pt-HARE5^Hs/Hs^* chimpanzee NPCs. n=1-2 cell lines of each condition; 5 individual differentiations; dots represents a cell line from a differentiation. **q**, Quantification of mitotic RGCs of D19 control and *Pt-HARE5^Hs/Hs^*chimp NPCs. n=1-2 cell lines of each condition from 3 individual differentiation; each grey dot represents an NPC island imaged and quantified; each color-coded dot represents the averaged value of the cell line from each differentiation. **r**, Experimental paradigm for cell cycle experiment in chimpanzee NPCs. **s**, Representative image of D19 control and *HARE5^Hs/Hs^* chimp NPCs stained with Ki67 (green), EdU (red) and DAPI (blue). **t**, Quantification of cell cycle exit of D19 control and *HARE5 ^Hs/Hs^* chimp NPCs. n=1-2 cell lines of each condition from 2 individual differentiation; Each grey dot represents an NPC island imaged and quantified; each dot represents a cell line from a differentiation. Scale bars: 50µm (h, l, s), 100µm (k, o). Graphs and bar plots, means ± S.D. *p<0.05, **p<0.01. One-way ANOVA with Dunnett’s multiple comparisons test (d), student’s unpaired, two-tailed t-test, comparison from colored dots (i, p), comparison from gray cell islands and images (j, m, q, t).

*HARE5* has four nucleotide changes discriminating humans and chimpanzees across the ultra-conserved region with *Hs-HARE5* showing two-fold higher enhancer activity than *Pt-HARE5* (Fig.5d). This begs the question of the extent to which each human-specific variant (I-IV) contributes to *Hs-HARE5* enhancer activity. To investigate this, we assayed *Pt*-*HARE5* luciferase reporters each containing only one human-specific variant (Fig.5c). Relative to the *Pt-HARE5* backbone, human variants I or II significantly increased activity by almost 60%. In comparison, variants III or IV showed only a 23% increase each (Fig.5d). Notably, *Pt-HARE5* carrying the first two human variants together (I+II), increased enhancer by ∼80% above the chimpanzee backbone, and comparable to *Hs-HARE5* (Fig.5d). This indicates that the first two variants have an especially strong role in promoting *Hs-HARE5* enhancer activity.

*De novo* copy-number variations (CNVs) and biallelic point mutations within HARs are associated with Autism Spectrum Disorder (ASD)^21,47^. Thus, we examined if there are any ASD-associated mutations in *Hs-HARE5* nearby these variants using databases^21^. Notably, 5 ASD mutations were reported in *HARE5*, including 2 substitutions and 3 zygosities^21^. In particular, an ASD-associated mutation (T>C) was located directly adjacent to site I in *Hs-HARE5* (Fig.5e). Using luciferase assays, we found this mutation significantly decreased *Hs-HARE5* enhancer activity (Fig.5f). Taken together, these experiments point to the functional importance of the *Hs-HARE5* locus for enhancer activity in human NPCs.

## CRISPR-edited human and chimpanzee NPCs reveal *Hs-HARE5* promotes proliferation

Having established human NPCs as a model for studying *Hs-HARE5*, we next tested the function of *Hs-HARE5* for human cortical development. For this, we generated *Pt-HARE5* knock-in human ESCs (H9 cells) using CRISPR/Cas9 genome editing (Fig.5g, Extended Data Fig.8a-c). Karyotyping, RT-qPCR, and OCT4 staining confirmed the knock-in cells were healthy and pluripotent (Extended Data Fig.8d).

We next used these lines to assess the functional impact of four chimpanzee nucleotides in *Hs-HARE5.* For this we generated human NPCs^48^ from 2 control *Hs-HARE5^Hs/Hs^*lines, including a parental H9 cell line and non-edited CRISPR line, as well as from 3 independent *Hs-HARE5^Pt/Pt^* lines (Fig.5g; Extended Data Fig.8f). To validate the forebrain cortical identity, we confirmed upregulation of *FOXG1* and *PAX6,* and downregulation of *OCT4* in D10-12 NPCs by RT-qPCR (Extended Data Fig.8h-j). Importantly, *FZD8* mRNA levels showed a significant reduction in D12-14 *Hs-HARE5^Pt/Pt^* NPCs compared to controls (Fig.5i), consistent with the function of this enhancer in promoting *FZD8* expression. As expected, we observed no effect upon *CCNY* levels (Extended Data Fig.8k). Thus, chimpanzee *HARE5* knock-in within human cells reduces expression of its target gene. This is consistent with reduced enhancer activity of *Pt-HARE5* observed in mice and human cells (Fig. 5d)^6^.

We then tested whether these four *HARE5* chimpanzee nucleotides impact human neurogenesis. Our mouse experiments revealed that *Hs-HARE5* promoted mitosis of early-stage RGCs (Fig3.d-e). We thus predicted that if this is true in human cells, replacing *Hs-HARE5* with *Pt-HARE5* would reduce human NPC proliferation. Indeed, quantification of SOX2+PH3+ cells showed a significant reduction in mitosis in the *Hs-HARE5^Pt/Pt^* NPCs (Fig.5h,j). This finding demonstrates that *Hs-HARE5* is essential for promoting proliferation of NPCs, consistent with observations in mice.

Next, we measured how human *HARE5* impacts human neurogenesis in cortical organoids which have more diverse cell composition and gene expression^49,50^. Towards this we utilized 2 control lines and 2 *Hs-HARE5^Pt/Pt^* lines (Extended Data Fig.8p) ^51^. To assess the impact on RGCs, we collected day 30 (D30) cortical organoids to model early developmental stages (Fig.5k, Extended Data Fig.8q). *FOXG1* RT-qPCR confirmed the forebrain cortical identity of these organoids (Extended Data Fig.8r). Immunostained sections for SOX2 further demonstrated expected neural rosette structures (Fig.5k,l). Notably, compared with controls, *Hs-HARE5^Pt/Pt^* organoids contained significantly fewer RGCs (Fig.5m). This is consistent with the findings in 2D human NPCs. These experiments demonstrate that just 4 chimpanzee-specific variants within human cells reduces proliferative capacity of progenitors.

Finally, we pursued an orthogonal experiment by modifying the *Pt-HARE5* locus in chimpanzee cells. To investigate how human *HARE5* substitutions impact chimpanzee neurogenesis, we generated *Hs-HARE5* knock in chimpanzee iPSCs by replacing the *Pt-HARE5* by CRISPR/Cas9^52^. We then generated chimpanzee NPCs from 2 control *Pt-HARE5^Pt/Pt^* lines including parental C3649 cells and non-edited CRISPR line, as well as 1 modified *Pt-HARE5^Hs/Hs^* line (Fig.5n; Extended Data Fig.8e) ^14,15^. *FOXG1* and *PAX6* were upregulated in D15 NPCs while *OCT4* was downregulated, confirming the forebrain identity (Extended Data Fig.8l-n). Notably, RT-qPCR showed higher *FZD8* mRNA levels in D15 *Pt-HARE5^Hs/Hs^*NPCs compared to controls (Fig.5p). We also observed a significant increase of *CCNY* levels in chimpanzee NPCs (Extended Data Fig.8o), which could suggest the chimpanzee NPC cellular environment differentially impacts target gene regulation by *Hs-HARE5*. Taken together, analysis of knock-in cell lines from two species show *HARE5* regulates *FZD8* expression in human and chimpanzee NPCs.

We then assessed proliferation of D19 NPCs using SOX2 and PH3 staining (Fig.5o). This revealed increased mitosis in the *Pt-HARE5^Hs/Hs^* NPCs (Fig.5q), indicating that *Hs-HARE5* promotes human progenitor proliferation. To further assess how *Hs-HARE5* impacts chimpanzee NPC proliferation, we performed a 24h EdU pulse (Fig.5r). Quantification of EdU and Ki67 revealed significantly fewer cells had exited the cell cycle (EdU+Ki67−/EdU+) in the *Pt-HARE5^Hs/Hs^* NPCs (Fig.5s,t). Thus, the presence of just 4 human-specific variants within chimpanzee cells is sufficient to enhance self-renewal and mitosis of progenitors. These findings are consistent with the increased mitosis and self-renewal of RGCs at the onset of neurogenesis in E12.5 *Hs-HARE5* mice.

Taken together, these findings demonstrate that *Hs-HARE5* functions as an RGC specific enhancer in the developing cortex. Strikingly, just four nucleotide changes in *Hs-HARE5* are sufficient to modulate its enhancer activity, which then promotes progenitor self-renewal and subsequent neural differentiation. These molecular, cellular, and developmental mechanisms underlie human features of cortical expansion (Fig.6).

**Fig. 6.**
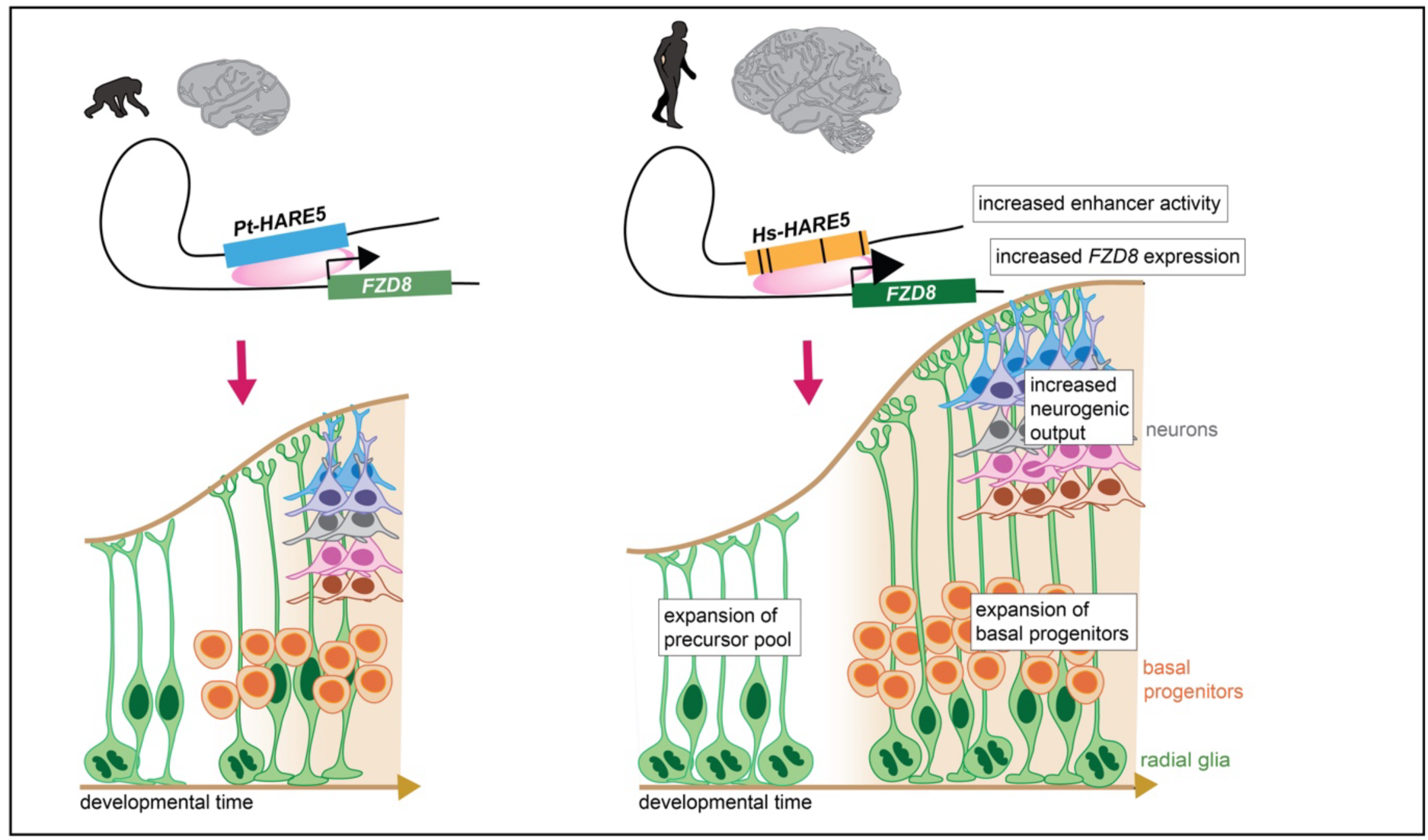
Summary findings of this study. *Hs-HARE5* includes 4 substitutions between human and chimpanzee. These 4 mutations contribute differently to the enhancer activity to drive the different expression level *FZD8.*In the developing cortex, *Hs-HARE5* activation is required for RGC proliferation at an early stage and promoted neurogenic differentiation later to produce more neurons in the cortex.

## Discussion

A long-standing question in biology is what molecular features make us uniquely human? Our genome contains thousands of human-specific DNA sequences^16,53,54^, including almost 3,000 Human Accelerated Regions (HARs), first computationally identified in 2006^4^. However, the *in vivo* function of HARs has been elusive and no study to date has shown that HARs directly impact human-specific brain features. Here, we use orthogonal models across three different species to demonstrate direct roles of a HAR in modifying critical events of human brain development. Specifically, we discover that the human *HARE5* enhancer contributes to cortical expansion and function by fine-tuning the behavior of radial glia, essential neural stem cells of the brain. Our study suggests that cortical evolution depends upon fine-tuned modification of key developmental pathways by gene regulatory changes.

We define developmental mechanisms by which *Hs-HARE5* promotes cortical expansion. Strikingly, *HARE5* and its target gene, *FZD8*, are predominantly active and expressed within RGCs. This is reflected by enhancer activity in mouse and human NPCs, as well as *FZD8* expression in the mouse, macaque, and human cortex. At the onset of neurogenesis, *Hs-HARE5* promotes RGC expansion by increasing self-renewal. This cellular modification aligns with the Radial Unit Hypothesis which proposes that increased founder cell number determines the size of cortical surface^55^. These findings are also consistent with studies of human and gorilla organoids showing that cortical expansion is influenced by transition and expansion of early stage progenitors^56^. As neurogenesis proceeds, *Hs-HARE5* accelerates differentiation of RGCs to preferentially produce IPs and neurons. How does *Hs-HARE5* influence the proliferative and neurogenic capacity of RGCs? We use live imaging, lineage analysis, scRNA-seq and transcriptomic analysis which pinpoints changes in RGC division mode and target gene expression. Together with amplified RGC number and capacity, increased abundance of BPs, including IPs and oRG/bRGs, seen in *Hs-HARE5* brains, contributes to neuronal expansion and enlarged brain size. Thus, *Hs-HARE5* promotes these human features by founder cell (RGCs) amplification, and production of basal progenitors and neurons across development.

*Hs-HARE5* promotes RGC proliferative and neurogenic capacity by amplifying *Frizzled8* expression in these precursors. This evolutionary modification impacts WNT signaling which is essential for neocortical development and implicated in disease^25,57^. The canonical WNT-signaling pathway, which depends upon β-catenin stabilization, primarily promotes RGC self-renewal^27,58,59^. In comparison, non-canonical WNT signaling is more linked to neural differentiation of NPCs^26,28,45,46^. Thus, our finding that *Hs-HARE5* inhibits canonical signaling at E14.5 correlates with the developmental impact of this enhancer from promoting RGC self-renewal to differentiation. We speculate that, due to its central role in development, during human evolution, the WNT signaling pathway was co-opted to modulate features of human brains. In future studies it will be fascinating to consider how other evolutionary modifications to components of this pathway synergize with *HARE5*. Beyond HARs, this may include HAQERs another recently discovered hominin-unique enhancers^53^.

What are the benefits of human-specific features such as increased cortical size on brain function and behavior? Although this question remains largely unanswered, changes in cortical architecture are likely to modify brain function in various ways^60,61^. For example, it may improve signal-to-noise ratio (SNR) and sensitivity, allowing for the extraction of more subtle features from sensory input. Indeed, increased cortical connectivity in mice that carry the human-specific gene *SRGAP2C* display improved sensory learning in a sensory discrimination task^62^. Increased neuron number may also permit additional processing pathways underlying the encoding and decoding of a wider range of behavioral variables or reflect an increase in storage capacity. Extracting more complex behavioral variables and having a larger amount of information available in memory from which to construct these variables could provide a basis for both improved learning and cognitive flexibility as it allows for generating novel behavioral solutions. Our data suggests that increased cortical neuron number in *Hs-HARE5* brains is associated with more independent functional activity, which is in line with an earlier study showing that increased cortical neuron number leads to more functional neuron ensembles and functional segregation of local neural networks^35^. Whether these changes in cortical activity provide a basis for increased memory storage, increased functional parcellation depending on behavioral demand, or improved cognitive flexibility ^63^ will be intriguing questions for future studies into how evolutionary changes in brain development modified brain function to facilitate improved learning and cognition.

Our study provides experimental paradigms for testing the most interesting candidate HARs and HAQERS. We previously use transgenic *Hs-HARE5-mFzd8* mice to implicate this locus in brain size due to accelerated cell cycle, however this model relied on overexpression of the *Fzd8* target gene^6^. With our new knock-in mouse model in which the *Mm-HARE5* locus is replaced with *Hs-HARE5* sequence, we can now pinpoint mechanistic developmental changes directly induced by the *Hs-HARE5* sequence itself. Importantly, we couple our findings with CRISPR-edited knock-in models of both human and chimpanzee cells.

The *HARE5* locus itself has notable modifications across evolution and linked with disease. In particular, *HARE5* has a segmental duplication (SD) present in humans, chimpanzees and gorilla^64^. Importantly, the SD regions are intact in our CRISPR edited cell lines. As this duplication presumably occurred before *HARE5* nucleotide substitutions, it would be interesting in future studies to consider how it affects primate features. Interestingly, the four substitutions in *HARE5* are highly conserved between modern humans and ancient humans (Denisova and Neanderthals)^65,66^. This suggests that brain size is relevant to *HARE5* specific substitutions, but that *HARE5* also impacts other features, related to the evolution of brain function. Further, the HARE5 locus also contains rare *de novo* mutations implicated in ASD^21,47,67^. We demonstrate that one ASD mutation, located adjacent to a human-specific variant, tempers enhancer activity in human NPCs. This fits with the notion that molecular and anatomical modifications in evolution are substrates for disease mutations^68^. Interestingly, both the human-specific variant 1 and ASD variant are predicted to influence the same transcription factor binding site^69^. Future studies will be valuable to understand the upstream transcriptional mechanisms modulating *HARE5* activity.

In sum, our study broadens our conceptual understanding of brain evolution, by demonstrating molecular, cellular and connectivity changes driven by human-specific gene regulation. This finding provides tangible support for the hypothesis proposed by King and Wilson that non-coding regulatory sequences account for biological differences between humans and chimpanzees^70^.

## Methods

### Mice

All mice experiments were approved by Duke IACUC and followed the guidelines from the Division of Laboratory Animal Resources from Duke University School of Medicine. Plug dates were defined as embryonic day (E) 0.5 on the morning the plug was identified. Emx1-Cre^34^and Tbr2-EGFP^40^ lines were previously described. *Mm-HARE5^Hs/Mm^* and *Mm-HARE5^Hs/Hs^* knock in mice were generated from Duke Mouse facility by using standard gene targeting techniques on 129/B6N hybrid mouse ES cells, the coordination of targeted mouse genome loci is (mm9 *chr18: 8871182-8872410*), the human *HARE5* sequence coordination is (hg38 *chr10:35949193-35950411*). Genotyping primers specific to the mouse lines are listed in SupplementaryTable1. All mice used in this study were from F8 or later generations on C57BL/6J background.

### DNA constructs

All oligos used for cloning are listed in SupplementaryTable1. All cloning PCR products were amplified by Q5 polymerase (NEB, M049L). For construction of vectors used for luciferase assays, human, chimpanzee, and mouse *HARE5* fragments were amplified from genomic DNA of individual species and inserted into the pNL1.1 vector (Promega) by KpnI and EcoRV enzyme sites. The sequences of the PCR primers and synthetic oligonucleotides are listed in SupplementaryTable1. For generating mutations *HARE5*-I, II, III, IV, V and ASD, we introduced mutations by primers and aligned by HiFi DNA Assembly Master Mix (NEB, E2621L). Ligated PCR products inserted into the pNL1.1 vector (Promega) by KpnI and EcoRV enzyme sites. Sequence was validated by Sanger sequencing. For CRISPR-Cas9 knock in of *Pt-HARE5* in Human ESCs and *Hs-HARE5* in Chimp iPSCs, guide RNAs were designed using the online tool from the lab of Feng Zhang (http://crispor.tefor.net/), we designed 2 pairs of guide RNAs which targeting both 5’ and 3’ side of *HARE5* loci (*Hs-HARE5* hg38 *chr10:35949493-35950111*; *Pt-HARE5* panTro4 *chr10:36445861-36446479*), referred as GuideL and GuideR. For cloning of the guides, briefly, the sense and antisense strand oligos for GuideL and GuideR were annealed and phosphorylated, and the duplexes were cloned into PX330 (Addgene, #158973) by BbsI site. Colonies were sequence validated using the U6_F primer. To increase the targeting efficiency, we amplified the whole sequence of U6_promoter till gRNA structure of GuideR and inserted it to GuideL vector to generate plasmid PX330-GuideL+R by BamHI enzyme site. For construction of Donor vectors used for CRISPR, human, chimpanzee *HARE5* fragments were amplified from genomic DNA of individual species, left and right homology arms (∼800bp) were amplified from genomic DNA of individual species with PAM site mutated. PCR fragments were ligased by HiFi DNA Assembly Master Mix (NEB, E2621L) and into the pCAG-Puro vector by XhoI enzyme site. Sequence was validated by Sanger sequencing.

### RNA extraction and RT-qPCR

For the mouse brain sample, embryonic cortices were dissected in cold PBS and RNA was purified from Trizol (Thermo, 15596026) method. For NPCs and organoids, RNA was purified using RNeasy micro-plus kit (Qiagen,74034). For mouse sample, 5ng cDNA was used for each qPCR reaction. For NPC and organoids characterizing, 15ng cDNA was used for each qPCR reaction. Primers were listed in SupplementaryTable1.

### Lentivirus packaging and transduction

HEK293T cultured in DMEM+10%FBS+P/S were used for lentivirus production. 26.75ug Transfer vectors, including pCAG dCas9-KRAB-2A-EGFP (addgene,92396) and gRNA-mcherry, 20ug Packaging plasmid (pCMV-deltaR8.9 or psPAX2) and 6.25ug Envelope plasmid (pCMV-VSVg or pMD2.G) were transfect with PEI-MAX (Polysciences, 24765-100) when cells reached to 70% in 15cm dish. Virus was harvest and concentrated 48hrs later. Human Neuroblastoma Cell Line SH-SY5Y cultured in DMEM+10%FBS+P/S were used for lentivirus transduction. Cells were collecting 3 days after for RNA extraction.

### Single cell RNA sequencing and analysis

E14.5 cortices were dissected in cold PBS, and dissociated using 0.25% Trypsin (Thermo, 25200056). Cell viability and number were counted by a Cell Counter (Thermo). The scRNA-seq experiment aimed to recover 8,000–10,000 cells from each embryo. cDNA amplification and library construction were performed following 10x Genomics Chromium Single Cell 3’ v3.1 kit. cDNA libraries were quantified using the Agilent High Sensitivity DNA Kit (Agilent, 5067-4626) and were sequenced on an Illumina NovaSeq. Samples were sequenced to a depth of 40,000-70,000 reads per cell. A total number of 22,695 cells from two control samples and 21,211 cells from two *Mm-HARE5^Hs/Hs^* sample were used for analysis, with an average of 2,110 genes detected per cell.

The 10x Genomics cell ranger count command version 6.1.2 was used to process fastq files and generate filtered count matrices. The count matrices were further processed using the Seurat R package version 4.0.5^71^. Normalization was performed using the SCTransform function, with the number of variable features = 5000, and regression of mitochondrial genes. The datasets were integrated using the IntegrateData function. A PCA was obtained with the RunPCA function with the number of principal components = 300. A UMAP was generated with the RunUMAP function based on the top 20 principal components. Clusters were obtained using the integrated counts with the FindNeighbors function, based on the 20 principal components. The function FindClusters was used with resolution = 1.1. Marker genes were obtained using the FindConservedMarkers function. The percentage of counts derived from MALAT1 transcripts were calculated, as high percentages are indicative of damaged neurons. Cells were considered damaged and removed if >5% of their counts were derived from MALAT1.

Cell cycle states were assigned with a classification approach using the cyclone function from the scran R package version 1.22.1^72,73^. This approach classifies each cell based on a reference dataset from mouse cells. It assigns each cell to either G1, G2/M, or S phases based on marker gene similarity in the reference dataset.

### Bulk RNA sequencing

For the mouse brain sample, E12.5 and E14.5 control and *Mm-HARE5^Hs/Hs^*cortices were dissected in cold PBS and RNA was purified from Trizol method, each genotype has 3 biological duplicates. cDNA libraries were prepared by Illumina TruSeq stranded mRNA kit. RNAseq libraries were sequenced on the NovaSeq (PE100) with 20M paired end reads. RNAseq libraries were sequenced to a depth of ∼40 million total reads per sample. RNASeq data quantified by Star Salmon from FASTAq files, gene expression was compared from E12.5 to E14.5 in both control and *Mm-HARE5^Hs/Hs^* genotypes individually. Differential gene expression (DGE) analysis using DESeq2. DGE lists were defined using an FDR <0.05, with Log2(FC)>=0.5 or <=-0.5. Overlapped and specific gene expression changes were defined by Biovinn (https://www.biovenn.nl/). GO analysis categories were selected by FDR <0.05, Plot were made by R Script.

### *FZD8* expression analysis

Processed bulk RNA-seq data was obtained from previous study^30^. Each data point represents a measurement in RPKM taken from a separate sample from a distinct neocortical area (DFC, ITC, MFC, OFC, STC, VFC, A1C, IPC, M1C, S1C, V1C). Human N=34, n=460. Macaque N=26, n=366. Human samples outside of the macaque age range (as measured by predicted equivalent human day) were excluded for statistical analysis.

### *In utero* electroporation (IUE)

*In utero* electroporation was performed as previously described^74^. Briefly, each E14.5 embryo was injected with 1ul of plasmid mix (containing 0.01% fast green and 500ng of pCAGGS-EGFP) and electroporation parameters: five 50 ms-pulses at 45V with 950 ms pulse-interval by platinum-plated BTX Tweezer Rodes. Plasmids were produced using Zymo Endotoxin Free Maxi Prep kits (Zymo, D4202) by following the manufacturers’ instructions.

### Cell Cycle exit

BrdU (Sigma, B5002) was dissolved in PBS for 10mg/mL stock. For cell cycle exit, BrdU was administered by IP injection at 50 mg/kg to pregnant dams at pregnant E11.5 or E13.5, embryos were harvested exactly 24 hr later at E12.5 or E14.5. BrdU were detected by antibody staining. For in vitro chimpanzee NPCs cell cycle exit, EdU was dissolved in PBS for 10mg/mL stock. 20µM EdU were added in the neural induction medium at D18, cells were harvested exactly 24 hr later at D19. EdU (Thermo, A10044) were detected by Click-iT® Plus EdU Imaging Kits (Thermo, C10638).

### Primary cultures and live imaging

Primary cortical cell culture was performed as described^39,41^. Briefly, Tbr2-EGFP positive embryos were pre-selected under fluorescent scope,microdissected both E14.5 dorsal cortices from each embryo and saved tails for genotyping. Cortices were trypsinized with 0.25% Trypsin for 6 min to create a single cell suspension, after centrifuge and suspension 100,000 cells were plated on poly-D-lysine(Sigma,P7280)-coated glass-bottom 24-well culture plates (MatTek, P24G-1.5-10-F). Cells were maintained in incubation chamber at 37°C and 5% CO2 for at least 2 hrs before moving to scope. Images were captured by 20x objective lens every 20 min with low exposure time for EGFP (30ms) and bright field (50ms) for 20-24hrs. Cell division were identified as previously^39^, and cell fate determination was performed post-imaging by immunostaining for Tuj1, and endogenous Tbr2-EGFP.

### Immunofluorescence staining and image acquisition

For histological analysis, samples (including mouse embryos, organoids, and NPCs) were collected and dissected in cold phosphate buffer saline (PBS), postnatal 21 and adult animals were performed with trans cardiac perfusion by PBS before dissection. Brains were fixed by immersion in a 4% Paraformaldehyde/PBS solution, the fixation time varied due to samples (overnight for embryonic and adult brains at 4℃, 3hrs for organoids at 4℃, 15mins for NPCs at room temperature). Following fixation, embryonic brains were cryoprotection overnight in 30% sucrose/PBS solution. Coronal cryosections were collected using a cryostat with a variable thickness depending on experiments (20mm for of embryonic immunofluorescence and in situ hybridization, 30mm for analysis of cortical layering at post-natal stages). Immunofluorescence was performed as described previously. Briefly, slides were permeabilized with a 10min wash in 0.25 % Triton-X/PBS. After blocking sections in 5% NGS/PBS for 1hr at room temperature. Primary antibody incubation in PBS overnight at 4℃ and secondary antibody for primary incubation for 1hr at room temperature. Antibody staining was listed below: NeuN (Cell Signaling,24307S,1/300); Pax6 (Millipore, AB2237,1/500); Sox2(Thermo,14-9811-82,1/1000); NeuroD2 (Cell Signaling,24307S,1/300); PH3 (Millipore, 06-570,1/1000); pVimentin-ser55(MBL,D076-3S,1/500); pVimentin-ser82 (MBL, D095-3, 1/500); Tbr2 (Abcam,ab183991,1/1000); Tuj1 (Covance,MM5-435P,1/1500); BrdU (Abcam,ab6326,1/200, need boil citrate 30min before blocking); Ki67 (Cell signaling,12202,1/300)

### Image analysis and quantification

Cell counting of embryonic brains and NPC cell cycle exit was performed in FIJI (ImageJ), using the Cell counter plugin. Cell counting of postnatal brains was done by Qupath automatically cell count. Cell counting of NPC cell mitosis island was done by Qupath automatically cell count combined with ImageJ. Cell counting of organoids was done by Qupath automatically cell count. Raw quantification data were initially processed by Excel (Microsoft) and then followed analyzed and graphed with Prism (GraphPad Software).

For binning analyses, X-Y coordinates were extracted from Cell counter data, and a script created in R was used to assign positive cells to specific bins, using the coordinates of the ventricular and pial borders as references. Bin numbers were reported in an Excel spreadsheet for analysis.

### iPSC, NPC, and organoid experiments

#### Knock in cell line generation

For establishment of the *Hs-HARE5^Pt/Pt^* mutant lines, plasmids PX330-hsGuideL+R (1µg) and pCAG-Puro-Pt-HARE5 (2µg) were electroporated into 1×10^6^ H9 cells using the P3 Primary Cell 4D-Nucleofector kit (Lonza, V4XP-3024). Following each electroporation, cells were separated to grow in 3 wells of a Matrigel (VWR, 354277) coated 6-well plate in mTesR^TM1^ supplemented with 1x CloneR (Stem Cell, 5888). After 3 days from electroporation, the cells were selected with 0.5 mg/ml Puromycin (Sigma, P8833). After approximately 5 days in selection medium, cells were switched back to mTesR^TM1^ medium until the single colony was big enough to passage. Single colonies were picked to Matrigel coated 24-well plates. After about 6 days, when the colonies reach ∼75% confluent in the well, colonies were split half for freezing and half for genotyping after dissociated by ReLeSR (Stem Cell, 5872). For the colonies genotyping, genomic DNA were extracted by Genomic DNA Purification Kit (Stem Cell, 79020) and firstly screened by PCR (Primer Pair1) and EcoRI digestion based on the third substitution gained EcoRI enzyme site in chimpanzee. The potential positive edited homozygote lines will be screened by a second round PCR (Primer Pair2) and Sanger sequencing to make the substitutions are correct. Primers listed in SupplementaryTable1. A total of 3 homozygote *Hs-HARE5^Pt/Pt^* lines were generated from ∼200 colonies. Homozygote colonies were tested Karyotypes through hPSC Genetic Analysis Kit (Stem Cell, 7550).

For establishment of the *Pt-HARE5^Hs/Hs^* lines, plasmids pX330-ptGuideL+R (1µg) and pCAG-Puro-Hs-HARE5 (2µg) were electroporated into 1×10^6^ C3649 cells using the P3 Primary Cell 4D-Nucleofector kit (Lonza, V4XP-3024). The selection and genotyping process was the same as above. A total of 1 homozygote *Pt-HARE5^Hs/Hs^* lines were generated from ∼150 colonies. Pluripotency was tested through OCT4 immunofluorescent staining.

#### Cell lines

All lines tested negative for mycoplasma contamination in this study were checked by the MycoAlert Mycoplasma Detection Kit (Lonza, LT07-518). One human ESC line WA09 (H9), one human iPSC line (8799), one chimpanzee iPSC line (C3649) were used in this study. H9 (WA09) was purchased from WiCell. 8799 was used in this study was previously characterized and genotyped^75^. C3649 was a gift from Yoav Gilad^52^. ESCs and iPSC lines were supplemented with mTesR^TM1^ (Stem Cell,85850) with Normacin (Invivogen, ant-nr-1). SH-SY5Y immortalized cell lines were maintained with DMEM-F12 media (Thermo Fisher, 11330057) supplemented with 10% fetal bovine serum (HyClone, SH30071.03HI) and 1% Pen-Strep (Thermo, 15140122). All cells were cultured at 37°C in 5% CO2 environment.

#### NPC differentiation

All ESCs (including H9, *Hs-HARE5^Hs/Hs^* control, *Hs-HARE5^Pt/Pt^-1*, *Hs-HARE5^Pt/Pt^-2*, *Hs-HARE5^Pt/Pt^-3*) and iPSCs (including 8799, C3649, *Pt-HARE5^Pt/Pt^* control, *Pt-HARE5^Hs/Hs^-1*) were cultured in Matrigel-coated plates with mTeSR^1TM^ media (Stem Cell,85850) in an undifferentiated state. Cells were passaged at a 1:5 ratio by treatment with ReLeSR (Stem Cell, 5872)/Accutase (Thermo, A11105) for maintaining.

For luciferase assay experiment, 8799 iPSCs were differentiated to neural progenitor cells (NPCs) as previously described^48^. Briefly, 0.8×10^6^ cells were seeded on Matrigel coated 60mm plates until reaching 100% confluent (Day 0), medium was replaced with neural induction medium containing 1 μM Dorsomorphin (Sigma, P5499), 10μM SB431542 (Selleckchem, S1067) every day. On Day 6, cells were washed with DMEMF12, dissociated with 1mg/mL DispaseII (Thermo, 17105041), clumps suspend in neural maintenance medium and plated on Poly-O-Ornithine (Sigma, P4957) /Laminin (Sigma, L2020) coated 6cm dishes. After D8, add 20ng/mL bFGF2 (R&D,233FB025) in neural maintenance medium when neural rosettes appeared.

For human cell NPCs experiment, H9, *Hs-HARE5^Hs/H^*^s^ control, *Hs-HARE5^Pt/Pt^-1*, *Hs-HARE5^Pt/Pt^-2*, *Hs-HARE5^Pt/Pt^-3* cells were differentiated to neural progenitor cells (NPCs) as above with minor changes^48^. Briefly, 0.8×10^6^ cells were seeded on Matrigel coated 60mm plates until reaching 100% confluent (Day 0), medium was replaced with neural induction medium containing 1μM Dorsomorphin (Sigma, P5499), 10μM SB431542 (Selleckchem, S1067) every day. At Day8-10, cells were washed with DMEMF12, dissociated with 1mg/mL DispaseII (Thermo, 17105041), clumps suspend in neural maintenance medium and plated on Poly-O-Ornithine/Lamini-coated 60mm dishes. After D10-12, add 20ng/mL bFGF2 in neural maintenance medium when neural rosettes appear. On Day12-14, human NPCs were washed with DMEMF12, dissociated with Accutase, and plated 2.5×10^^5^ cells on Poly-O-Ornithine/Lamini-coated 4-well chamber slides (Mattek, CCS-4) for 2days for further IF and analysis.

For chimpanzee NPCs experiment, C3649, *Pt-HARE5^Pt/Pt^* control and *Pt-HARE5^Hs/Hs^ −1* cells were differentiated to neural progenitor cells (NPCs) as described^14^. Briefly, 0.7×10^5^ cells were seeded on Matrigel coated 12-well plate (Day 0), medium was replaced with N2B27 neural induction medium containing 50 ng/mL bFGF2(R&D,233FB025), 100 ng/mL Noggin (Miltenyi Biotec,130-103-456) every day. At Day15, chimp NPCs were washed with DMEMF12, dissociated with Accutase and plated 2×10^^5^ cells on Poly-O-Ornithine/Lamini-coated 4-well chamber slides (Mattek, CCS-4) for 4days for further IF and analysis.

#### Cortical organoid generation

Forebrain organoids were generated as described^50^. Briefly, on day 0, H9, *Hs-HARE5^Hs/Hs^* control, *Hs-HARE5^Pt/Pt^-1*, *Hs-HARE5^Pt/Pt^-2* cells were dissociated to single cells by Accutase, with 9,000 cells per well were reaggregated in ultra-low cell-adhesion 96-well plates with V-bottomed conical wells (S-bio, MS-9096VZ) in Cortical Differentiation Medium (CDM) I. Day 0-6, ROCK inhibitor Y-27632(Stem Cell, 72304) was added (final concentration of 20 mM). Day 0-18, Wnt inhibitor IWR1 (Millipore, 681669) and SB43154 (Selleckchem, S1067) were added (final concentration of 3 and 5 mM, respectively). From day 18, the aggregates were cultured in ultra-low attachment culture dishes under orbital agitation in CDM II until day 30.

#### Luciferase assays

Human NPCs differentiated from iPSC line (8799), cells were split and plate at D10 by Accutase. 4×10^5^ cells were plating on 24-well plate coated with Poly-O-Ornithine/Lamini-coated. On day14, cells were transfected using Lipofectamine LTX reagent (Thermo Fisher Scientific, A12621) the following day with 250ng either mouse or chimpanzee and human mutation pNL1.1 luciferase vectors generated with enhancer sequences as described above. 250ng Firefly plasmid was co-transfected to control transfection efficiency. The luciferase assays were performed 24 h after transfection using the Nano-Glo Dual-Luciferase Reporter Assay System (Promega, N1551) according to the manufacturer’s instructions. Luciferase activity was measured and quantified by Luminescence Microplate Reader System. For each experiment, the raw value was normalized by pNL1.1 empty vector. Data were collected from at least 3 independent differentiations.

### Widefield calcium imaging in vivo

#### Mouse surgery

Adult mice (>P65) were used for functional imaging experiments. Age-matched wild-type littermates, as well as C57Bl6 mice obtained from Jax, were used as controls. All mice were maintained on a 12h light/dark cycle at 20-22°C and 30-70% humidity. Adult mice were anesthetized using isoflurane, after which they were injected with adeno-associated virus (AAV-PHP.eB-syn-jGCaMP7s, Addgene) using a 30-gauge needle. Mice were subsequently returned to the home cage to allow for brain-wide neuronal expression of GCaMP7s. Three weeks after virus injection, mice were surgically implanted with a custom designed headplate for widefield imaging. Mice were anesthetized with 1-2% isoflurane, injected with meloxicam subcutaneously (5 mg/kg body weight), and placed in a stereotactic frame (Neurostar). A medial incision was made, and the skin was retracted, exposing the skull over the dorsal cortex. The exposed skull was cleaned, and three layers of cyanoacrylate glue (Zap-A-Gap CA+, Pacer Technology) were applied to clear the bone. After the glue was dry, a custom-designed titanium headplate was secured onto the skull using Metabond dental acrylic (Parkell) to allow for head-restraint during widefield calcium imaging.

#### Widefield imaging

Resting state neural dynamics were captured using widefield calcium imaging of GCaMP fluorescence ^76^. The exposed dorsal cortex was illuminated using two consecutively strobing LEDs. A blue LED (Thorlabs, M470L5) with bandpass filter (Semrock, FF01-460/60-25) was used to excited GCaMP fluorescence, while a green LED (Thorlabs, M530L3) and bandpass filter Semrock, FF01-530/43-25) was used to captured total hemoglobin absorption. A Semrock bandpass filter (FF01-565/133-25) was placed in front of the camera to eliminate the GCaMP excitation light. Near-simultaneous imaging of total hemoglobin the green LED was necessary to remove hemodynamic signals that are known to confound GCaMP data^76^. GCaMP fluorescence was recorded using an Andor Zyla 4.2+ sCMOS camera and Nikon lens (Nikon Micro-Nikkor 60 mm f/2.8D) at 60Hz. Resting state imaging, in the absence of any external stimulus, was done consecutively for 10 minutes.

#### Image processing and analysis

To analyze widefield neural activity, the hemodynamic confound was first removed from the GCaMP signal using green reflectance as previously described^76^. We then performed principal component analysis (PCA) to denoise the data and for each mouse, the number of components needed to account for 85% of their respective total variance was kept. Following this, the data were spatiotemporally clustered in an unsupervised manner using k-means clustering (correlation-based with 120 components, 60 per hemisphere). K-means clustering was used to ensure that any changes to functional parcellation due to increased neuron number in mice expressing *Hs-HARE5* could be detected. To determine the correlation value threshold of independence, we calculated Pearson’s correlation between each pixel’s neural time course and the average time courses of each k-means cluster. The correlation threshold of independence (max r between regions) is the maximum value at which a given pixel is correlated to any cluster beyond its own. Thus, neural activity within a given pixel is considered independent at max r because it does not have a stronger correlation with any other functional region other than the cluster it belongs to. To determine the within-region correlation, Pearson’s correlation analysis was done on each pixel relative to only the cluster to which they belong. Finally, to contextualize the functional clustering and correlation analyses, we grouped each k-means cluster into known anatomical cortical regions based on the Allen Atlas Common Coordinate Framework. The correlation threshold of independence was then averaged for all regions within a given Allen Atlas CCF region.

## Statistical analysis

Mouse samples were selected unbiased for various experimental conditions, samples were excluded from severely damaged or dead, or if the immunostaining signal or IUE efficiency was clearly lower than normal or was not comparable between the different groups of samples. Primary cultured mouse NPCs were excluded from cell death either during dissection or after imaging. Mouse experiment was performed from at least 2 independent litters with at least 3 animals for each condition, the sample sizes, biological and duplications are listed in the figure legends.

Human/Chimpanzee NPCs and cortical organoids samples were selected based on the RT-qPCR of *FOXG1/PAX6/OCT4* levels, samples were excluded based on the extremely not comparable *FOXG1* level from each differentiation. All analyses were performed blindly by 1 or more investigators. No predetermination of sample sizes was carried out because our research is an exploratory study. The sample sizes are listed in the figure legends. The number of data points, biological replicates and statistical tests used for all the comparisons are indicated in the figure legends. Human and chimpanzee cells experiments were performed from at least 2 independent differentiations, the sample sizes, biological and duplications are listed in the figure legends.

The parametric tests used were two-tailed unpaired or paired Student’s t-test; one-way analysis of variance (ANOVA) followed by Dunnett’s multiple comparisons test; or two-way ANOVA followed by Bonferroni’s post-hoc test; Chi-square test. The statistical test used for each experiment was specified in the figure legends.

## Acknowledgements

We thank Silver lab members, Craig Lowe for helpful discussions. We are grateful for Nonhuman primate biological materials provided by the Emory National Primate Research Center (EPC) f/k/a Yerkes National Primate Research Center; the Chimpanzee iPSC line was a gift from Dr. Yoav Gilad. We thank the Duke Cancer Institute for generating *Hs-HARE5* knock-in mice, Duke Mouse Transgenic Facility for mouse maintenance, Duke light and microscopy core for use of shared equipment, Duke Sequencing Core for scRNA-seq library sequencing. The following funding supported this research: D.L.S. (NIH: R01NS083897, R01NS120667, R01NS110388, R01MH132089-02), J.L. (Ruth K. Broad fellowship), Emory National Primate Research Center Grant (No. ORIP/OD P51OD011132).

## Author contributions

J.L. and D.L.S. conceived of and designed the study. J.L. and D.L.S. wrote the manuscript. J.L, M.L, Y.M., C.F.E. performed experiments and analyses of the mouse model. J.E.S.F., A.J.M., G.W. performed scRNA-seq library preparation and analysis. H.M.D. and A.M.S. performed *FZD8* expression analysis across primates. H.Z and E.S. performed adult neural connectivity experiments. F.M. and J.L. performed *HARE5* mutation cloning and luciferase experiments. F.M., C.M.M and J.L. performed CRISPR editing on human and chimpanzee ES/iPSCs, F.M. and J.L. performed cortical organoid experiments with the help of C.F.E. for data analysis. J.L. performed human and chimpanzee NPC differentiation experiments with the help of C.F.E. for data analysis. All authors reviewed, edited, and approved the manuscript.

## Competing interests

The authors declare no competing interests.

**Correspondence and requests for materials** should be addressed to Debra Silver.

## Data and materials availability

Datasets used in this study will be publicly available at the NCBI Gene Expression Omnibus (GEO) database (www.ncbi.nlm.nih.gov/geo).

---

**Extended Data Fig. 1.**
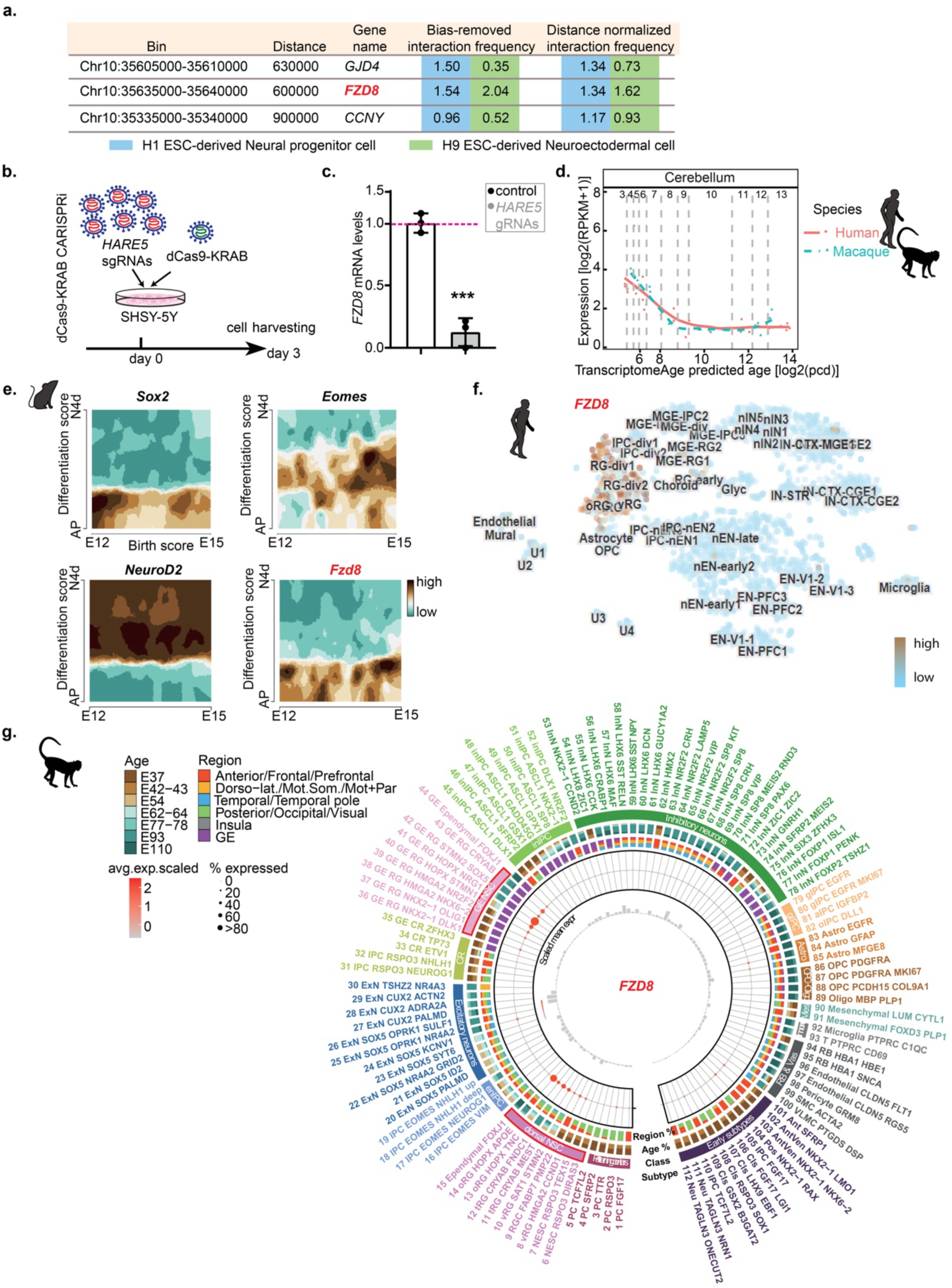
*FZD8* is the main target of *HARE5* and expressed in radial glia. **a**, HiC data indicated *FZD8* is the highest frequency target of *HARE5* in human neural cells. **b**, Experimental paradigm for CRIPSRi of *HARE5* by lentivirus in SH-SY5Y cell line. **c**, RT-qPCR of *FZD8* mRNA after 3 days transduced with control and *HARE5* gRNAs. n=3 individual transduction. **d**, Published bulk RNA-sequencing analyses confirm *FZD8* expression is comparable in cerebellar region across human and macaque. **e-g,** Published single-cell RNA-sequencing analyses confirm *Fzd8* expression is restricted to RGCs in mice cortex from E12-E15(**e**), and in human (**f**)and macaque (**g**)fetal cortices at peak neurogenesis stages. Graphs and bar plots, means ± S.D. ****p<0.0001. Student’s unpaired, two-tailed t-test (c).

**Extended Data Fig. 2.**
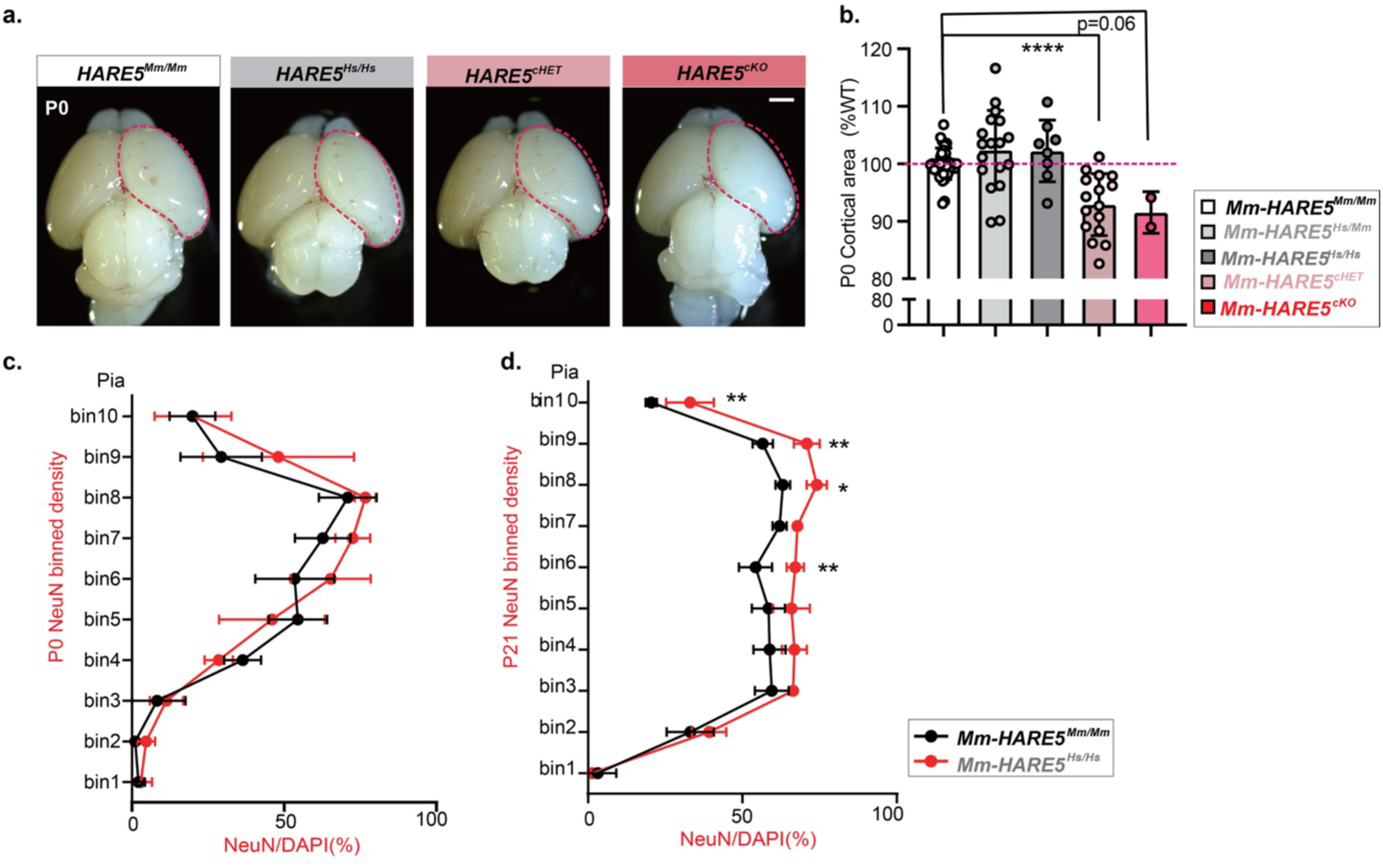
*HARE5* impact on cortical size and neuronal distribution. **a**, Representative whole mount images of control, *HARE5^Hs/Mm^, HARE5^Hs/Hs^, HARE5^cHET^* and *HARE5^cKO^* brains at P0. Dotted line is control superimposed on both brains. **b**, Quantification of cortical area at P0. Each litter normalized with wild type littermates; dots represent brains. **c**, Binned quantification of NeuN+/DAPI percentage within each bin in P0 control and *HARE5 ^Hs/Hs^* cortices. Bin 1 is apical lining the ventricle, and Bin 10 is at the pia. X-axis represents the percentage. n=3-7 animals; 5 litters per genotype. **d**, Binned quantification of NeuN+/DAPI percentage within each bin in P21control and *HARE5^Hs/Hs^* cortices. n=4-6 animals; 3 litters per genotype. Scale bars: 1mm (a). Graphs and bar plots, means ± S.D. *p<0.05, **p<0.01, ***p<0.001. ****p<0.0001. One-way ANOVA with Dunnett’s multiple comparisons test (b), two-way ANOVA with Bonferroni’s multiple comparisons test (c,d).

**Extended Data Fig. 3.**
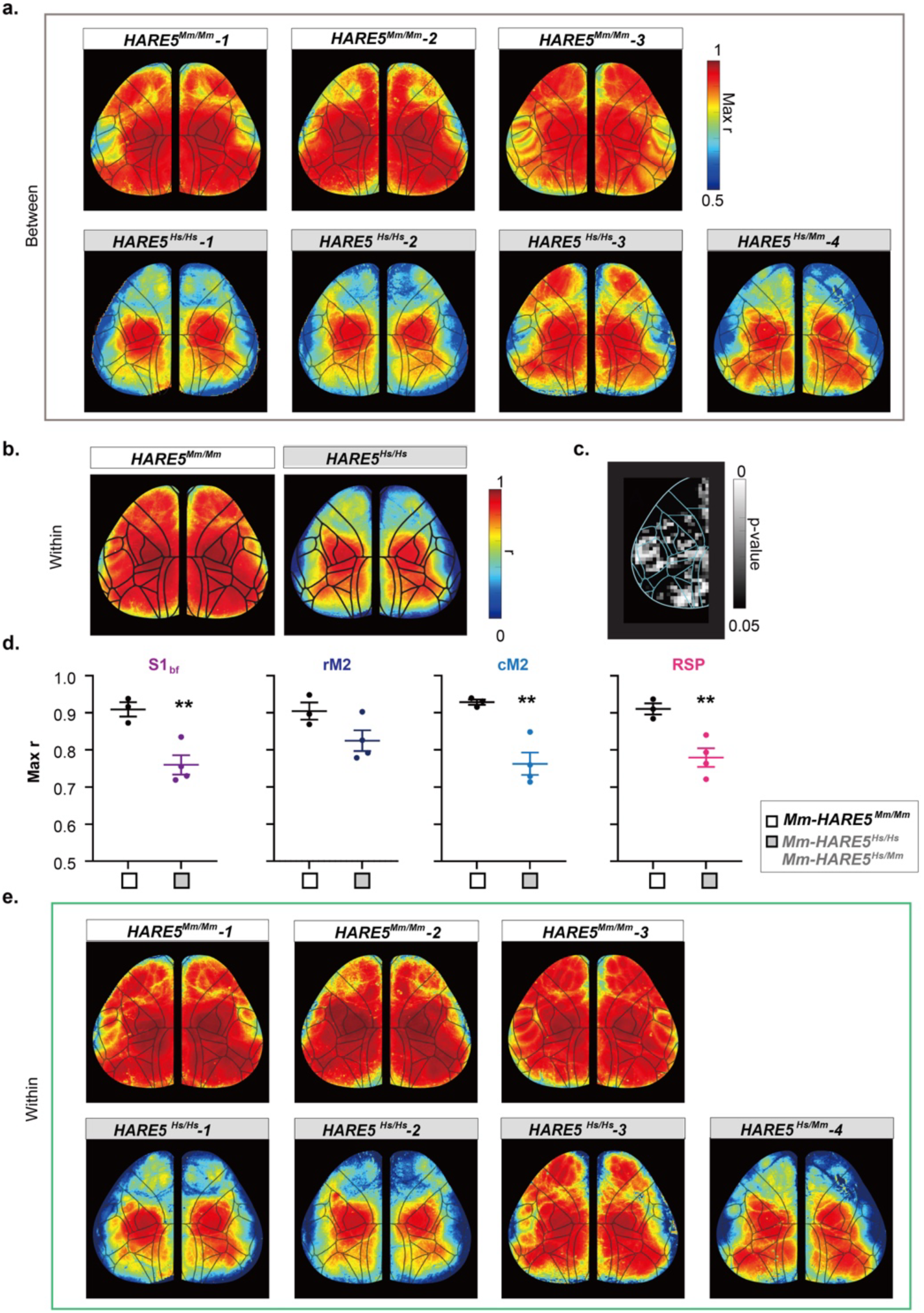
Correlation analysis of resting state activity of each individual mouse. **a**, Spatial maps show the pixelwise correlation value threshold of independence across the imaging field of view for each mouse. **b-e**, A Similar analysis as Fig.2. (**b**) Maps showing the temporal correlation between each pixel and the average time course of the functional cluster it belongs to. (**c**) Correlation maps were spatially binned into a grid of 10×10 pixels and compared between genotypes. P-value map shows statistically significant spatial grids as calculated using two-tailed t-test (All non-black pixels are statistically significant, p<0.05). (**d**) Functional clusters were grouped into known anatomical regions based on the Allen Atlas Common Coordinate Framework. Secondary motor cortex (M2) was split rostral-caudally and evaluated separately. Graphs show average Pearson’s correlation (r) within regions (whisker barrel field S1, rostral M2, caudal M2, and RSP). Each data point represents an individual animal. (**e**) Spatial maps show pixelwise correlation of each pixel to its own functional cluster (within region correlation) for each individual mouse. **h-j** was done to evaluate pixelwise correlation *within* functional clusters.Graphs and bar plot, means ± S.D. *p<0.05, **p<0.01, ***p<0.001. ns, not significant. Student’s unpaired, two-tailed t-test (d).

**Extended Data Fig. 4.**
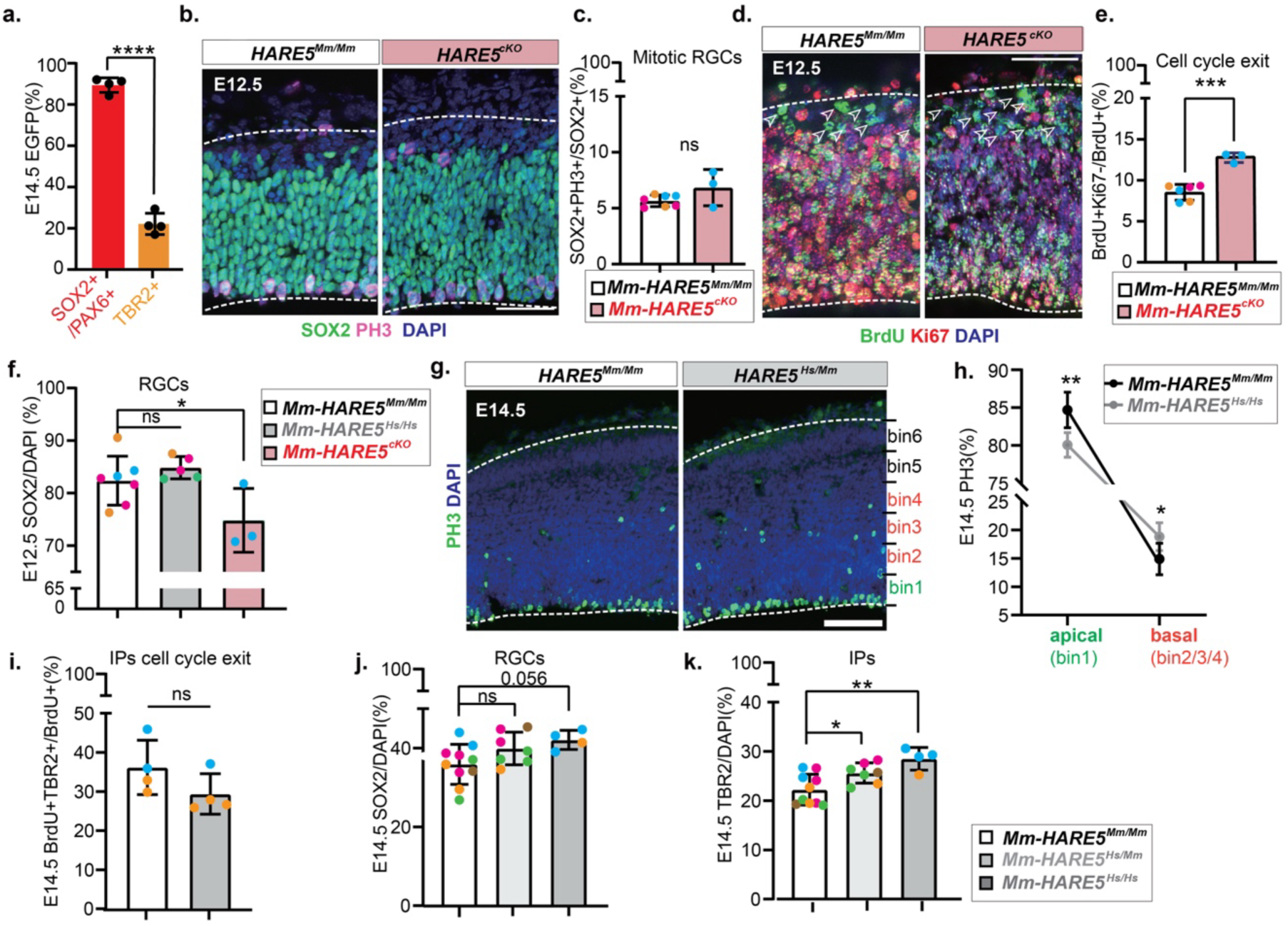
*HARE5* impact on RGC and IP proliferation. **a**, Quantification of proportion of RGCs (PAX6+ or SOX2+) and IPs (TBR2+) from EGFP+ cells in E14.5 *Hs-HARE5*::EGFP cortices. n=4 embryos; 3 litters. **b**, Representative sections of E12.5 control and *HARE5^cKO^* brains stained with SOX2 (green), PH3 (magenta) and DAPI (blue). **c**, Quantification of mitotic RGCs in E12.5 control and *HARE5^Hs/Hs^*cortices. n=3-7 embryos; 4 litters per genotype. **d**, Representative sections of E12.5 control, *HARE5^cKO^* brains stained with BrdU (green), Ki67 (red) and DAPI (blue). **e**, Quantification of cell cycle exit in E12.5 control and *HARE5^cKO^* cortices. n=3-6 embryos; 4 litters per genotype. **f**, Quantification of RGCs in E12.5 control, *HARE5^Hs/Hs^* and *HARE5^cKO^* cortices. n=3-7 embryos; 4 litters per genotype. **g**, Representative sections of E14.5 control and *HARE5^Hs/Mm^* brains stained with PH3 (green) and DAPI (blue). **h**, Quantification of PH3+ proportion in E14.5 control and *HARE5^Hs/Mm^* cortices. n=3-4 embryos;3 litters per genotype. **i**, Quantification of IPs cell cycle exit ratio in E14.5 control and *HARE5^Hs/Hs^* cortices. n=4 embryos ;2 litters per genotype. **j**, Quantification of SOX2+ RGCs in E14.5 control, *HARE5^Hs/Mm^*and *HARE5^Hs/Hs^* cortices. n=4-10 embryos; 5 litters per genotype. **k**, Quantification of TBR2+ IPs in E14.5 control, *HARE5^Hs/Mm^* and *HARE5^Hs/Hs^*cortices. n=4-10 embryos; 5 litters per genotype. Scale bars: 50µm (b, d, g). Graphs and bar plots, means ± S.D. *p<0.05, **p<0.01, ***p<0.001. ****p<0.0001. One-way ANOVA with Dunnett’s multiple comparisons test (f, j, k), student’s unpaired, two-tailed t-test (a, c, e, i), two-way ANOVA with Bonferroni’s multiple comparisons test (h).

**Extended Data Fig. 5.**
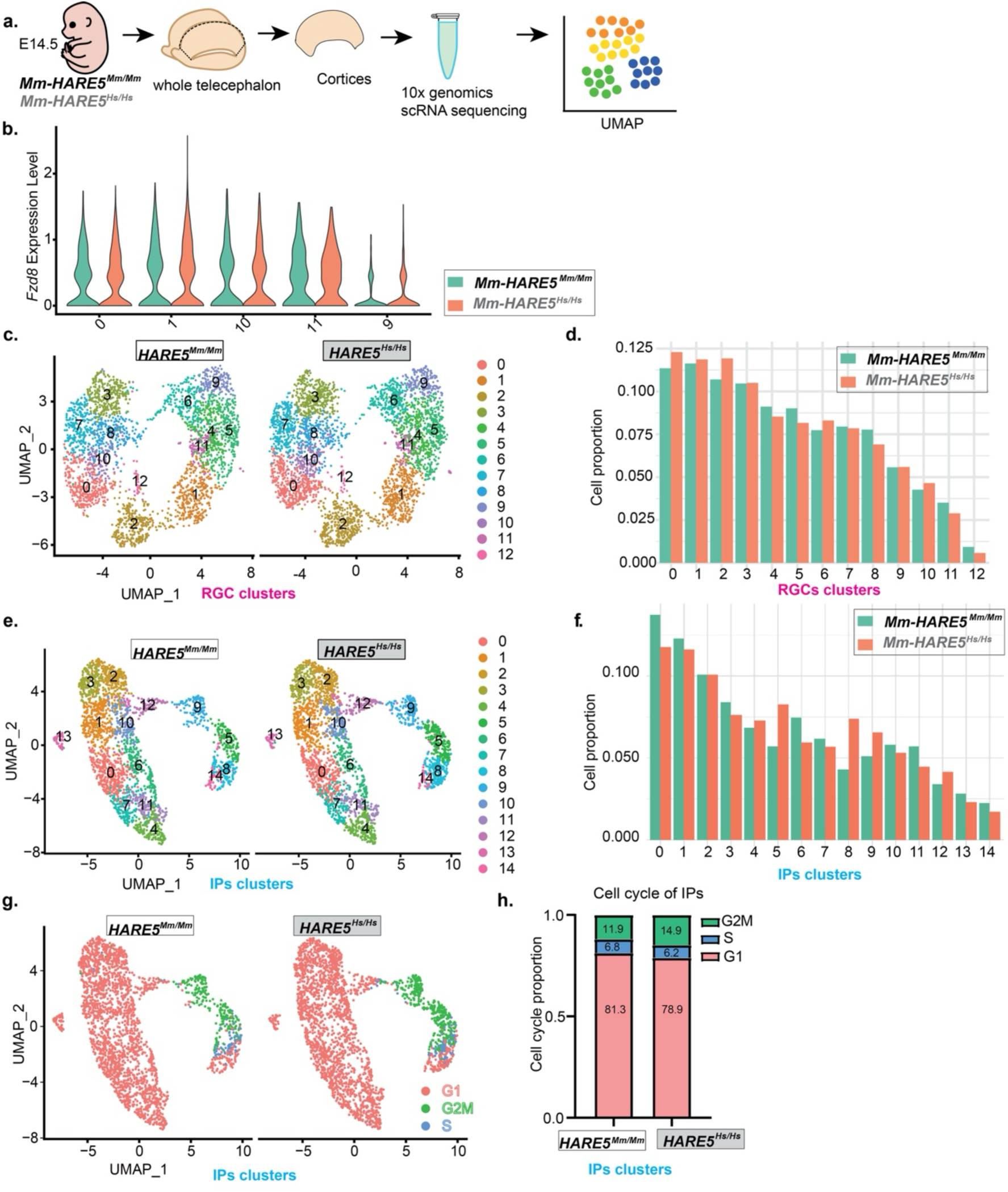
scRNA-seq indicates cell composition changes in *Hs-HARE5* cortices. **a**, Experimental paradigm for scRNA sequencing of E14.5 control and *HARE5^Hs/Hs^*cortex. **b**, Violin plots showed *Fzd8* highly expressed RGC clusters in E14.5 control and *HARE5^Hs/Hs^*cortex. n=2 embryos ;2 litters per genotype. **c,e**, UMAP of RGCs (**c**) and IPs (**e**) from 4 samples. Cell clusters were characterized by gene list from Supplementary Table 2. n=2 litters per genotype. **d,f**, Quantification of RGCs(**d**) and IPs (**f**) subclusters proportion. Cell clusters were characterized by gene list from Supplementary Table 3. n=2 embryos; 2 litters per genotype**. g**, UMAP of IPs cell cycle status from 4 samples. n=2 embryos; 2 litters per genotype. **h**, Quantification of G2M IPs in E14.5 control and *HARE5 ^Hs/Hs^*cortices by counts. n=2 embryos; 2 litters per genotype.

**Extended Data Fig. 6.**
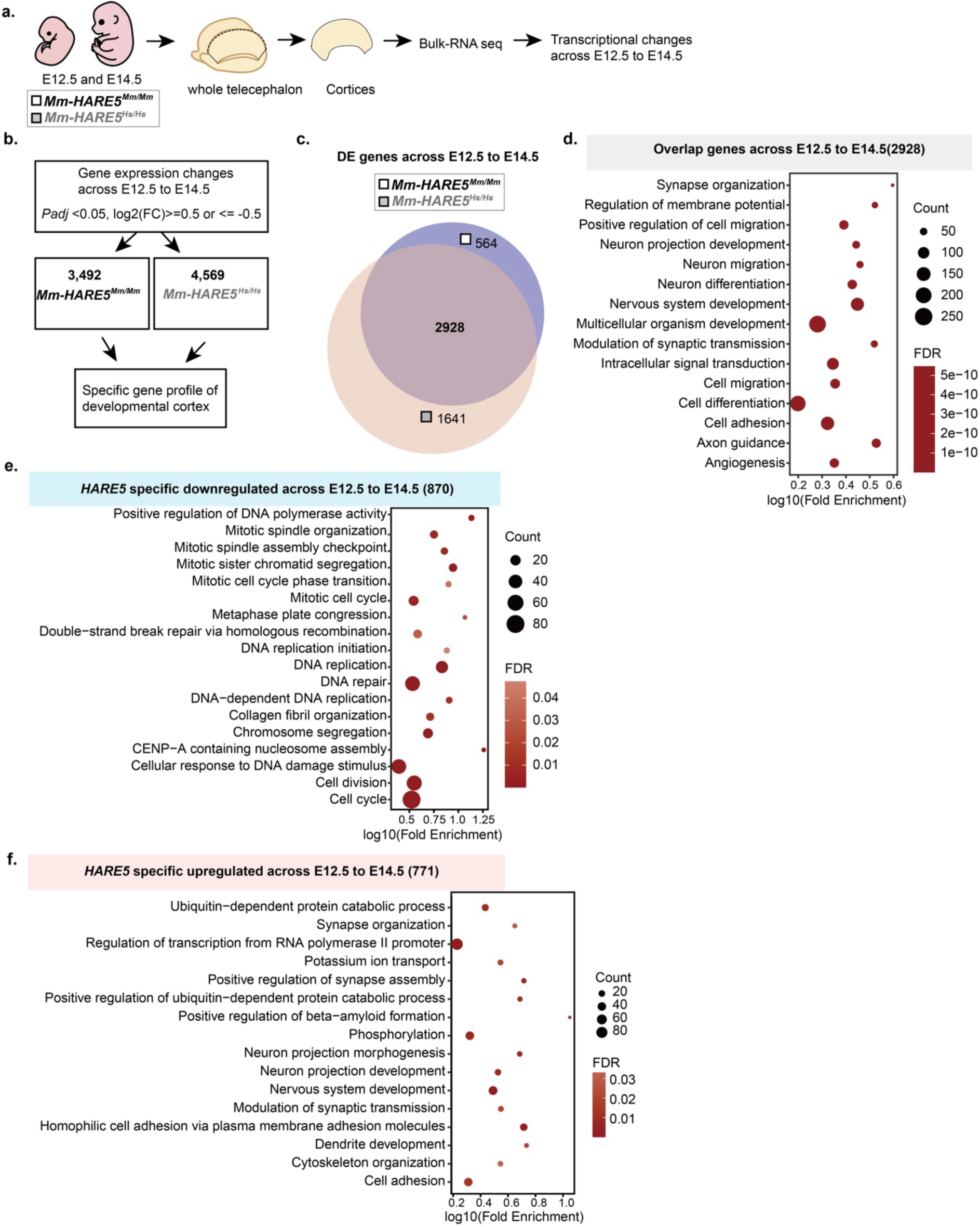
*Hs-HARE5*-dependent transcriptome is associated with neurogenic transition across cortical development. **a**, Scheme representing the sample collection for Bulk-RNA sequencing of control and *HARE5^Hs/Hs^* cortex. Cortices were collected at E12.5 and E14.5. n=3 embryos; 2-3 litters per genotype. **b**, Criteria used to identify statistically significant genes during cortical development across E12.5 to E14.5 in *HARE5^Hs/Hs^* cortices. **c**, Proportional Venn diagram of the genes expressed in controls, *HARE5^Hs/Hs^* cortices, and their overlaps. The diagram was drawn using BioVenn. DE genes are listed in Supplementary Table 4. **d**, Gene Ontology analysis showing overlap genes across cortical development from E12.5 to E14.5 in both genotypes. DE genes are listed in Supplementary Table 4. **e**, Gene Ontology analysis showing statistically significant enrichment for downregulated genes in *HARE5^Hs/Hs^* cortices. DE genes are listed in Supplementary Table 5. **f**, Gene Ontology analysis showing statistically significant enrichment for upregulated genes in *HARE5^Hs/Hs^* cortices. DE genes are listed in Supplementary Table 5.

**Extended Data Fig. 7.**
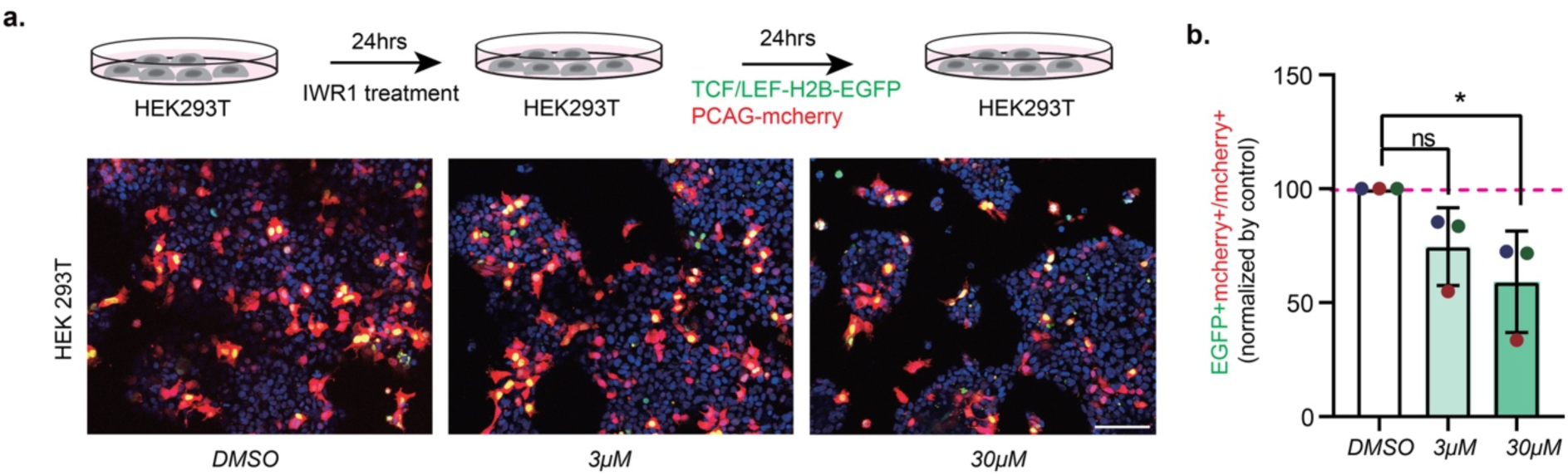
TCF1/Lef-H2B-EGFP reporter is sensitive to detect canonical WNT-signalling changes. **a**, Experimental paradigm for testing canonical WNT-signal by reporter after inhibition by IWR1 in HEK cell line. **b**, Quantification of WNT activated cell proportion in cells treated with DMSO, *3µM* IWR1 and *3µM* IWR1. n=3 independent transfection. Each dot represents the average value of different imaging field from each transfection. Scale bars: 100µm (a). Graphs and bar plots, means ± S.D. *p<0.05, **p<0.01, ***p<0.001. ****p<0.0001. One-way ANOVA with Dunnett’s multiple comparisons test (b).

**Extended Data Fig. 8.**
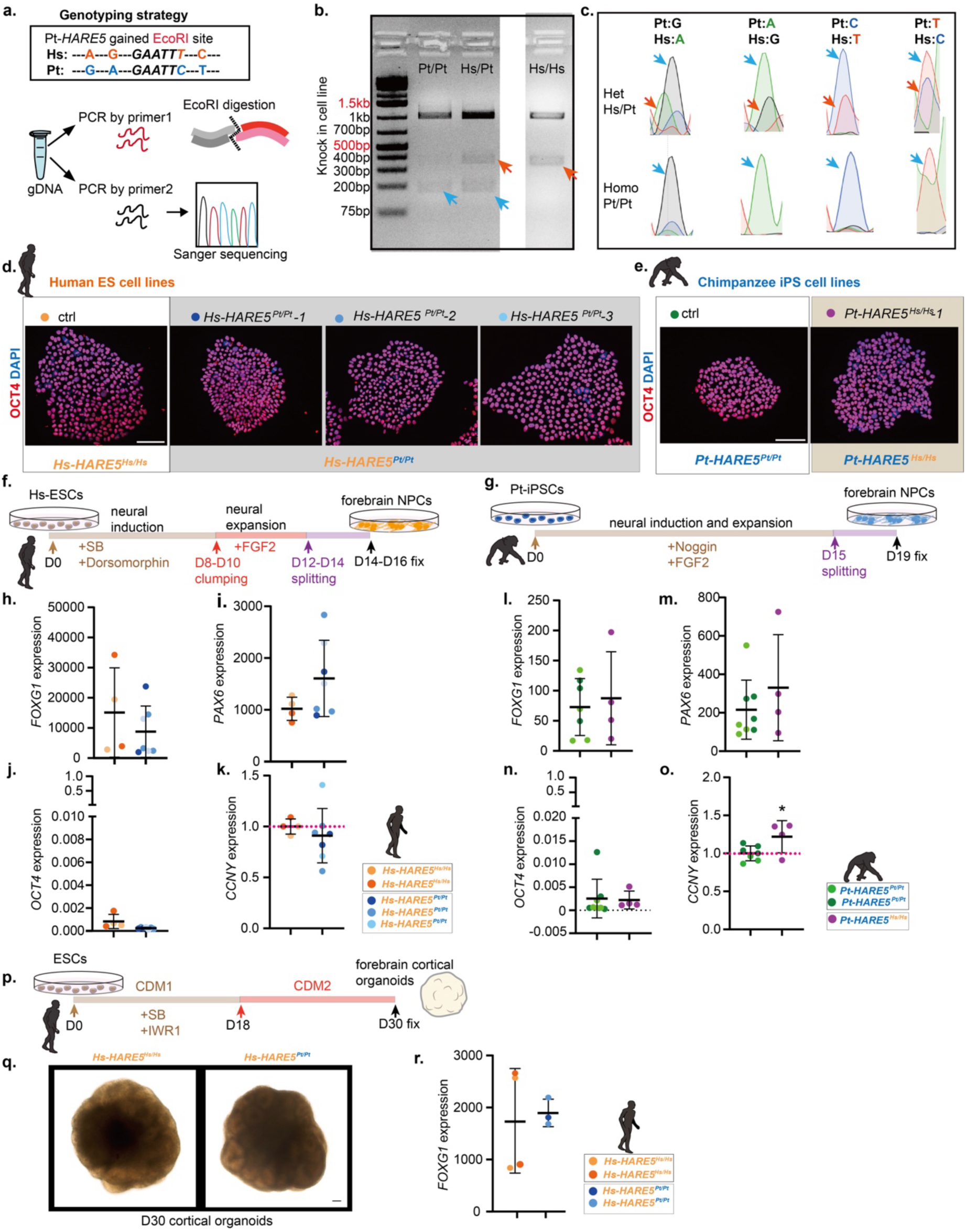
*HARE5* knock-in hiPSC/hESC lines and cortical NPCs/organoid differentiation. **a**, Experimental paradigm for genotyping after CRISPR editing. **b**, EcoRI enzyme digestion showed the successful mutation between *Hs-HARE5* and *Pt-HARE5* (gained EcoRI site within the 3^rd^ substitution) in the control and mutant cell line. **c**, Sanger sequencing showed the successful knock-in with all 4 mutations in human *Hs-HARE5^Pt/Pt^*lines compared with heterozygote *Hs-HARE5^Hs/Pt^* lines. **d,e**, Representative colonies of controls, *Hs-HARE5^Pt/Pt^* (**d**) and *Pt-HARE5^Hs/Hs^* (**e**) stained with OCT4 (red) and DAPI (blue). **f**, Experimental paradigm for human NPCs differentiation. **g**, Experimental paradigm for chimpanzee NPCs differentiation. **h-j**, Human NPC identity characterization by RT-qPCR on D12-14 of controls (2 lines) and *Hs-HARE5^Pt/Pt^* lines (3 lines). (**h**) qPCR of *FOXG1* mRNA; (**i**) qPCR of *PAX6* mRNA and (**j**) qPCR of *OCT4* mRNA. Normalized with D0 H9, n=2-3 individual differentiation shown as individual dots each. **k**, RT-qPCR of *CCNY* mRNA of D12-14 human NPCs of controls (2 lines) and *Hs-HARE5^Pt/Pt^* lines (3 lines). Normalized with controls, n=2-3 individual differentiation shown as individual dots each. **l-n**, Chimpanzee NPC identity characterization by RT-qPCR on D15 of controls (2 lines) and *Pt-HARE5^Hs/Hs^* lines (1 line). (**l**) qPCR of *FOXG1* mRNA; (**m**) qPCR of *PAX6* mRNA and (**n**) qPCR of *OCT4* mRNA. Normalized with D0 C3649, n=4 individual differentiation shown as individual dots each. **o**, RT-qPCR of *CCNY* mRNA of D15 chimp NPCs of controls (2 lines) and *Pt-HARE5 ^Hs/Hs^* lines (1 line). Normalized with controls, n=4 individual differentiation shown as individual dots each. **p**, Experimental paradigm for human 3D cortical organoids differentiation. **q**, D30 cortical organoids in bright field. **r**, RT-qPCR of *FOXG1* mRNA of D30 human organoids of controls (2 lines) and *Hs-HARE5 ^Pt/Pt^* lines (2 lines) normalized with D0 H9. n=1 individual differentiation, each dot represents individual organoid. Scale bars: 100µm (c,q). Graphs and bar plots, means ± S.D. *p<0.05, **p<0.01, ***p<0.001. ****p<0.0001. Student’s unpaired, two-tailed t-test (k,o).

## Notes

### Competing Interest Statement

The authors have declared no competing interest.

